# Myeloid STING restrains cardiac remodeling by suppressing macrophage amyloid precursor protein

**DOI:** 10.64898/2026.07.01.735895

**Authors:** Niranjana Natarajan, Ebin Johny, Varsha Sriram, Mika Hara, Praise Ama Antwi, Lee Ohayon-Steckel, Aarush Dutta, Adarsh Raj, Partha Dutta

## Abstract

Mitochondrial DNA (mtDNA) released into the cytosol activates innate immune signaling and promotes inflammation, yet its role in macrophages following sterile tissue injury remains poorly understood. Here, we show that cardiac macrophages from both patients and mice with myocardial infarction (MI) exhibit increased mitochondrial biogenesis, mitochondrial content, membrane potential, and expression of mitochondrial nucleases that facilitate mtDNA release. Consistently, macrophage-specific silencing of genes regulating mitochondrial biogenesis or mtDNA processing attenuated adverse cardiac remodeling after MI. Unexpectedly, despite the role of mtDNA in activating the cGAS–STING pathway, myeloid deletion or macrophage-specific silencing of *Sting* or *cGas* exacerbated ventricular dilation, fibrosis, and contractile dysfunction following MI. Single-cell transcriptomic and cell communication analyses identified amyloid precursor protein (APP) as a key downstream effector of STING in cardiac macrophages. Macrophage-specific *in vivo App* silencing rescued the detrimental effects of myeloid *Sting* deficiency, establishing APP as a critical mediator of adverse remodeling. Mechanistically, STING interacted with the transcriptional repressor MZF1, promoted its nuclear localization, facilitated its binding to the *App* promoter, and suppressed *App* transcription to restrain adverse cardiac remodeling. Together, our findings uncover an unexpected cardioprotective function of myeloid STING and identify the STING–MZF1–APP axis as a previously unrecognized mechanism governing cardiac repair after myocardial infarction.

## Introduction

The ability of macrophages to exert a wide range of functions ranging from tissue homeostasis to inflammatory response stems from their phenotypic plasticity and ability to adopt varied activation states based on extrinsic cues^1^. Different macrophage polarization states are associated with unique metabolic profiles and shift in mitochondrial metabolism, structure, and function^2^. In addition to modulating macrophage polarization and function, damaged mitochondria and elevated ROS levels result in the release of mtDNA in the cytosol, which acts as a damage associated molecular pattern (DAMP), mediating the type I interferon response via binding to cyclic GMP-AMP synthase (cGAS) and activating STimulator of Interferon Genes (STING) ^3,4^.

The cGAS-STING pathway is a crucial component of the innate immune system and serves to detect double stranded DNA (dsDNA) in the cytoplasm, independent of other DNA sensing pathways ^4,5^. cGAS binds to dsDNA and generates cyclic guanosine monophosphate adenosine monophosphate (cGAMP), a second messenger that activates STING, which in turn initiates the transcription of type-1 interferons ^3,4,6^. STING is expressed in a wide range of immune cell types - macrophages, dendritic cells, lymphocytes, and in non-immune cells like endothelial and epithelial cells ^7,8,9^. At steady state, STING localizes to the ER. Upon cGAMP binding, STING forms a complex with TANK-binding kinase 1 (TBK1) that translocates to the perinuclear region and activates transcription factors IRF3 and NF-kB by phosphorylation, resulting in the stimulation of type I interferon gene expression and cytokine production ^9^.

Various metabolic pathways, such as liver X receptor-mediated lipid metabolism ^10^ and glycolysis ^11^ can regulate the cGAS-STING pathway. Although cGAS primarily serves to detect pathogenic DNA in the cytoplasm, it is also activated by cytoplasmic nuclear and mitochondrial DNA ^3,4,12^. Pathological release of mtDNA into the cytosol via the mPTP and VDAC-dependent channels activates the cGAS-STING pathway and fuels inflammation ^13,14^ whereas mitophagy by stimulating PINK1 and Parkin can subside this effect ^15,16^. mtDNA-mediated uncontrolled inflammation is critical in aging-related neurodegeneration ^17^, multiple sclerosis ^18^, and platelet activation and granule secretion ^19^. Aberrant cGAS-STING signaling has also been implicated in autoimmune disease, cancer, and diabetes ^20^. STING agonists, when used in combination with chemotherapeutic agents, augmented their anti-tumor effects ^21,22^.

In cardiovascular diseases, chronic activation of STING has been associated with NF-kB and NLRP3 inflammasome signaling, leading to the production of IL-1B in diabetic cardiomyopathy and following ischemia reperfusion injury ^23–26^. Moreover, STING has been shown to aggravate cardiac remodeling after MI ^27,28^ and viral myocarditis ^29^. Global knockdown or inhibition of STING improved cardiac function and ameliorated cardiac hypertrophy in a murine model of DCM and decreased inflammation and apoptosis in cultured cardiomyocytes ^23,24^. In contrast to these studies revealing the deleterious functions of STING in cardiovascular study, a recent study reported a protective function of STING in a mouse model of transverse aortic constriction ^30^. These contrasting findings of STING functions in cardiovascular disease could be attributed to the lack of cell-specific deletion or targeting of this nucleic acid sensor. In support of this statement, STING manifests opposing functions in cancer depending on cell type by which it is expressed^21,31^. Furthermore, the role of myeloid STING in cardiovascular disease and heart failure is not understood.

The main goals of this study are two-fold: A) To understand how mitochondrial DNA synthesis in cardiac macrophages impacts STING and post-MI inflammation and B) To delineate the function of myeloid STING in cardiac remodeling following MI. We demonstrate that cardiac macrophages in the infarct heighten mitochondrial biogenesis and the expression of the proteins involved in this process, mtDNA release into the cytosol, and the expression of mitochondrial nucleases, which facilitate cleavage of mtDNA and its egress from mitochondria. Ablation of the genes driving mitochondrial biogenesis, such as *Pgc1a* and *Polg*, and mitochondrial nucleases, such as *Fen1* and *Apex2*, in macrophages subsided inflammation and cardiac fibrosis and remodeling after MI. These data indicate that the cGAS-STING pathway, which is activated in response to oxidized and fragmented mtDNA, is detrimental in post-MI healing of the heart. Surprisingly, genetic deletion and silencing of *Sting* and cGas in macrophages resulted in adverse cardiac remodeling and exaggerated fibrosis by inflating the production of APP after MI. Mechanistically, STING bound to the suppressor MZF1, facilitating its nuclear translocation. MZF1 bound to the *App* promoter, resulting in suppressed transcription of this gene.

## Results

### Cardiac macrophages after MI display increased mitochondrial volume and membrane potential

Oxidized and fragmented mitochondrial DNA exacerbates inflammation and contributes to disease pathogenesis ^32–35^. However, how mitochondrial biogenesis affects inflammation after an acute sterile injury, such as MI, is not known. In this study, we used a mouse model of MI and cardiac specimens from patients with a history of MI and ischemic cardiomyopathy to characterize mitochondria in cardiac macrophages, which are crucial in propagating inflammation and tissue remodeling. We observed that mitochondrial volume as assessed by TOM20 immunostaining and oxidative DNA damage indicated by 8-OHdG staining were elevated in cardiac macrophages of the patients with MI compared to healthy controls (**Figures 1A-1B**). Similarly, cardiac macrophages from mice with MI had elevated TOM20 and 8-OHdG immunostaining, indicating greater mitochondrial volume and increased oxidative DNA damage, respectively (**Figures 1C-1D**). Cardiac macrophages but not monocytes in the infarcts exhibited greater mitochondrial volume (**Figure 1E**) and mitochondrial membrane potential (**Figure 1F**). With the exception of bone marrow neutrophils and splenic monocytes, mitochondrial volume and membrane potential were unaltered after MI in most leukocytes and myeloid cell populations in the bone marrow and spleen (**Figure S1A-S1B**). Increased mitochondrial volume is primarily attributed to augmented mitochondrial biogenesis associated with mitochondrial DNA (mtDNA) synthesis. To examine if exaggerated inflammation, which is associated with post-MI sequelae, augments mtDNA synthesis, we treated bone marrow-derived macrophages (BMDM) with oxidized LDL (oxLDL). This treatment significantly augmented mitochondrial DNA content, assessed by D-loop/Tert ratio (**Figure 1G**). In line with this observation, oxLDL treatment upregulated the expression of several genes involved in nucleotide biosynthesis, such as *Adsl*, *Ctps2*, *Gmps,* and *Umps* (**Figure S1C**). We next sought to examine the association of circulating free mtDNA with cardiac function and systemic inflammation in patients with ST-elevation MI (STEMI). The circulating mtDNA levels were inversely correlated with left ventricular ejection fraction and were positively correlated with the inflammatory cytokine levels (**Figure 1H**). Altogether, these data indicate that MI escalates mtDNA synthesis in cardiac macrophages.

**Figure 1:**
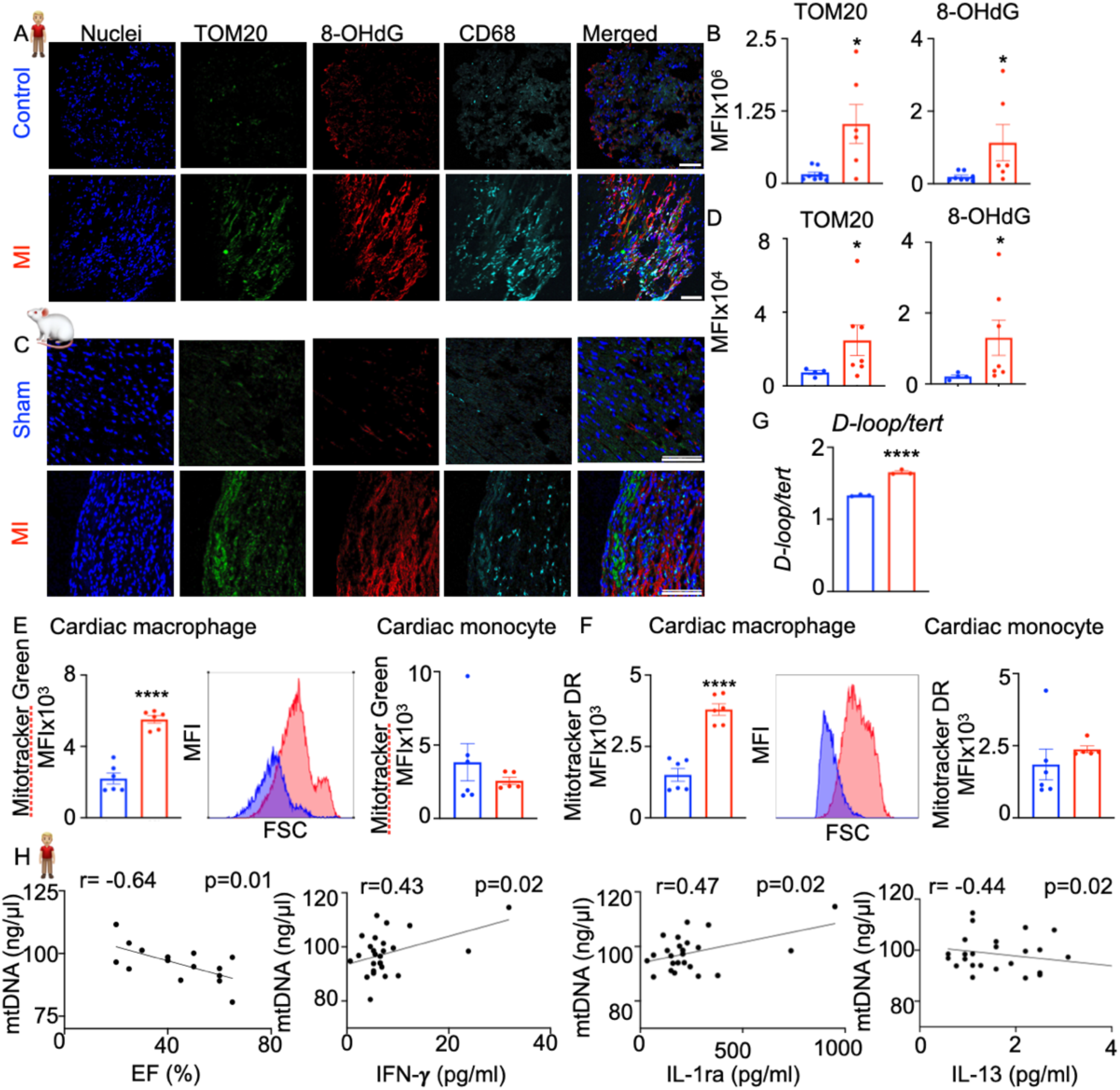
MI augments mitochondrial volume, membrane potential, and DNA synthesis. A- D) TOM20 (mitochondrial volume) and 8-OHdG (oxidative DNA damage) in cardiac macrophages of patients with a history of MI and matched controls (A and B) and mice with sham surgery and MI (C and D) were quantified by confocal microscopy. scale bar:100μm. n=8 (control), 6 (HF), 4 (sham), and 7 (MI). E-F) Mitochondrial volume (E) and membrane potential (F) in cardiac macrophages and monocytes of mice on day 7 after MI were assessed by flow cytometry. n=6/ group. G) D-loop/Tert ratios were calculated to measure mitochondrial DNA in native (n) and oxidized (ox) LDL treated macrophages, n=4. H) mtDNA was assessed and various cytokine levels were evaluated in the plasma of patients with STEMI. The correlations among mtDNA levels, cardiac ejection fraction (EF) (n=15), and the cytokine levels are depicted (n=26). EF data were available only in 15 patients. \**p*<0.05, \*\*\*\**p*<0.0001, two-tailed Student’s t-test, Pearson correlation analysis.

### Targeting macrophage mitochondrial biogenesis genes improves cardiac function after MI

It has been documented that new mitochondrial DNA synthesis aids NLRP3 inflammasome activation and type I interferon gene expression ^36^. The expression of CMPK2, POLG and PGC1A was greater in cardiac macrophages in MI hearts, compared to sham-operated control (**Figures 2A-2B**). Based on these data and the associations among circulating mtDNA levels, cardiac function, and inflammation, we hypothesized that aberrant mtDNA synthesis contributes to cardiac remodeling after MI. Towards this end, we silenced key genes involved in mitochondrial biogenesis – *Cmpk2, Polg,* and *Pgc1a*- specifically in macrophages using a lipidoid nanoparticle approach (**S2A**) ^32^. While the treatment did not show any significant improvement in cardiac function at 7 days after MI (**Figure S2B**), these mice had improved cardiac output and stroke volume compared to control at day 28 after MI (**Figure 2C-2D**). Although ejection fraction was improved in mice treated with *siPolg* and *siPgc1a,* diastolic and systolic volumes were unaltered. The abundance of Immune cell populations was not altered in the heart, spleen and bone marrow after the gene silencing (**Figure S2C-S2E**). Next, we examined the effects of *Cmpk2* and *Polg* on inflammatory gene expression in murine bone marrow-derived macrophages. *Cmpk2* genetic deficiency and *Polg* and *Pgc1a* silencing in macrophages displayed a muted inflammatory response in response to oxLDL stimulation (**Figure 2E**).

**Figure 2:**
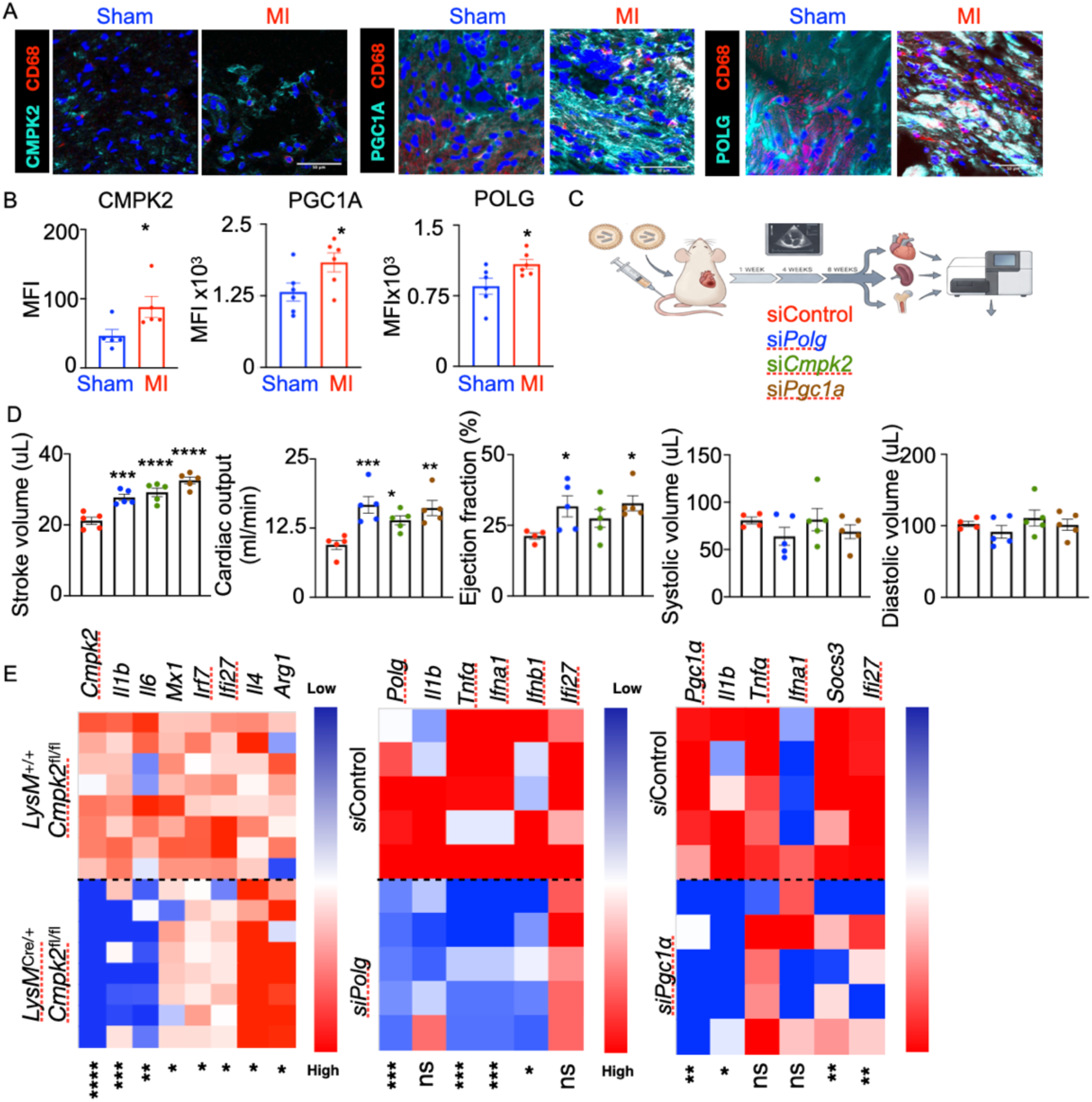
Macrophage-specific silencing of mitochondrial biogenesis genes improves post-MI cardiac function. A) Immunofluorescence staining was carried out to detect CMPK2, PGC1A, and POLG in the infarct 28-days post-MI, n=6. B) Experimental design to silence *Cmpk2*, *Pgc1a,* and *Polg* in a macrophage-specific manner after MI. C) A cartoon containing the experimental outline is shown. D) Stroke volume, cardiac output, ejection fraction, and systolic and diastolic volume on day 28 after MI are quantified, n=5. E) Inflammatory gene expression was analyzed in BMDM lacking *Cmpk2*, *Polg*, or *Pgc1a* using qPCR, n=5. \**p*<0.05, \*\**p*<0.01, \*\*\**p*<0.001, \*\*\*\**p*<0.0001, two-tailed Student’s t-test (A,E), one-way ANOVA with post-hoc Fisher LSD test (D).

### Mitochondrial nucleases FEN1 and APEX 2 increase inflammation and post-MI cardiac remodeling

Inflammatory macrophages exhibit greater oxidative DNA damage, as observed by higher 8-OHdG levels in cardiac macrophages in the infarct (**Figure 1A-D**). Damaged mitochondrial DNA is cleaved by the nucleases FEN1, MGME1, and APEX2 and released into the cytoplasm via VDAC, mPTP or BAK/BAX-dependent mechanisms ^14^ (**Figure 3A**). We first measured the expression of mitochondrial nucleases *Fen1*, *Mgme1*, *Apex2,* and *Exog* in cardiac macrophages isolated from sham and MI hearts and BMDM exposed to oxLDL. While the expression of *Fen1*, *Mgme1*, and *Apex2* was upregulated in BMDM after oxLDL treatment (**Figure 3B**), only *Fen1* and *Apex2* expression was elevated in cardiac macrophages following MI (**Figure 3C**). Congruently, the protein expression of these two nucleases was magnified in cardiac macrophages of mice (**Figure S3A**) and patients (**Figures 3D**) with MI. We observed exuberant cytoplasmic mtDNA in macrophages after oxLDL exposure by quantifying mitochondria (TOM20)-free oxidized (8-OHdG^+^) mtDNA (TFAM^+^) (**Figure S3B**), suggesting mtDNA egress in the cytosol in inflammatory conditions. In line with this finding, silencing the genes encoding the nucleases significantly downregulated the inflammatory response to oxLDL in BMDM (**Figure 3E**). Both nuclear and mtDNA (B-DNA) can supercoil to form Z-DNA, which can be detected by Z-DNA binding protein 1 (ZBP1), triggering cGAS-STING activation and type I interferon response ^37^. We observed that while total and mitochondrial Z-DNA and B-DNA levels were elevated in response to oxLDL, ZBP1 levels were unaltered (**Figure S4A-S4D**). Damaged and fragmented mtDNA can be released into the cytoplasm through VDAC, the mPTP or the BAX/BAK complex^14^. Congruently, VDAC inhibition and *Bak* and *Bax* silencing attenuated oxLDL-driven inflammatory response in BMDM (**Figure S5A-S5B**). The nucleic acid sensors *Ifih1*, *Ddx58* (RIG-I), and *Sting* detect double stranded DNA in the cytoplasm to stimulate downstream inflammatory responses. *siIfih1*, *siDdx58,* and *siSting* treatment lowered TNF-⍺ secretion in BMDM in response to oxLDL, *siIfih1* and *siDdx58* decreased IL-1β secretion, and *siSting* diminished IL-6 levels (**Figure S5C**).

**Figure 3:**
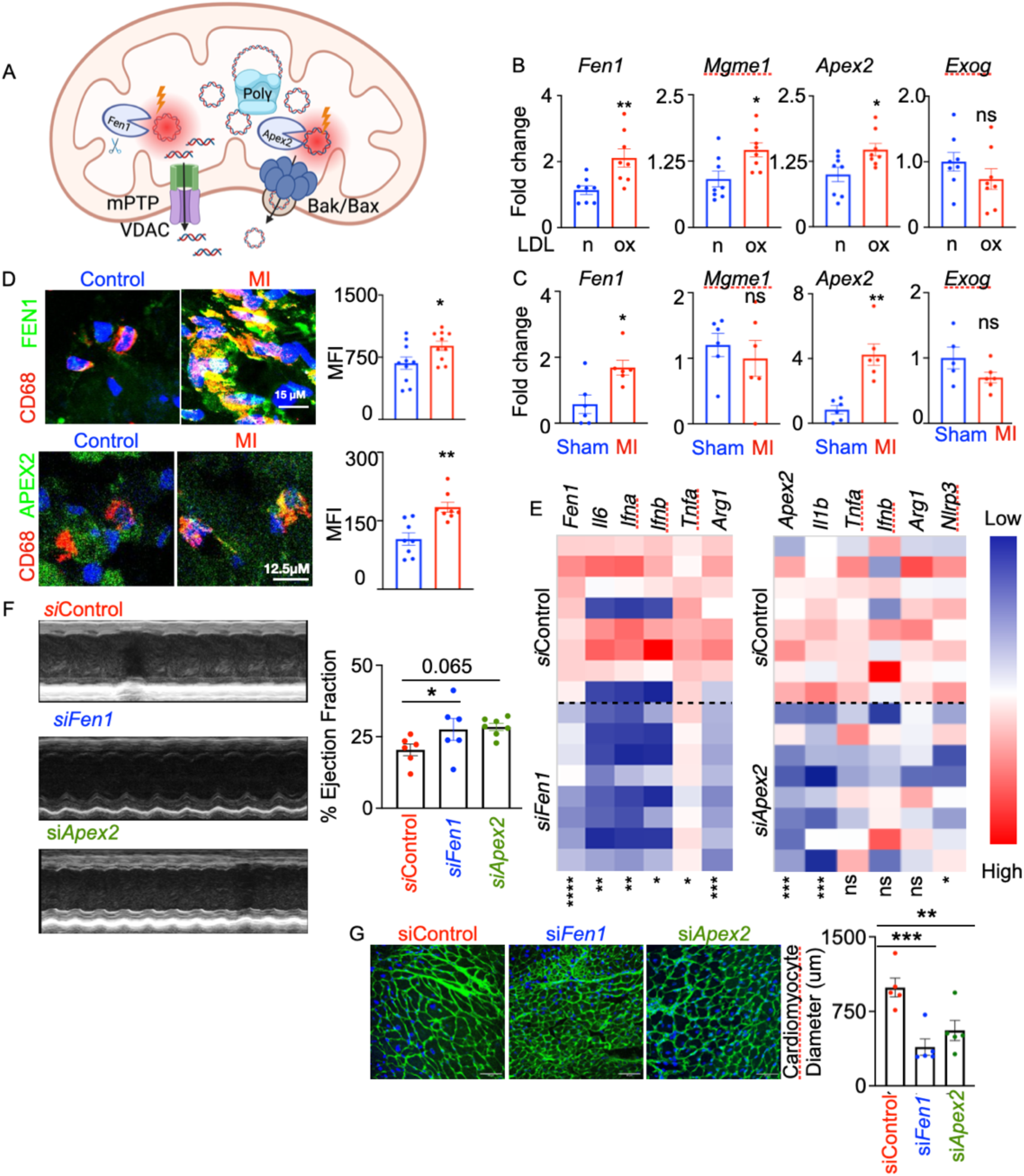
The mitochondrial nucleases FEN1 and APEX2 promote inflammation and post-MI cardiac remodeling. A) Schematic of mtDNA release into cytoplasm is provided. B-C) The expression of the mitochondrial nucleases in either nLDL or oxLDL-treated macrophages (B, n=8/ group) and cardiac macrophages from mice with sham surgery or MI (C, n=6). D) FEN1 and APEX2 MFI were quantified by confocal microscopy in the heart of patients with MI and control patients, n=10/ group, scale bar: 15μm (FEN1), 12.5μm (APEX2). E) Inflammatory gene expression in BMDM upon silencing of *Fen1* and *Apex2* was evaluated by qPCR (n=6/group). F) Mice were treated with either *siControl*, *siApex2* or si*Fen1* i.v for four weeks after MI. Echocardiography was performed in these mice to measure systolic function and left ventricular ejection fraction in these mice (n=5/group). G) Cardiomyocyte size in the mice with MI and siRNA treatment was discerned (n=5/group). \**p*<0.05, \*\**p*<0.01, \*\*\**p*<0.001, \*\*\*\**p*<0.0001, two-tailed Student’s t-test (B-E), one-way ANOVA with post-hoc Fisher LSD test (F,G).

Based on our observations that APEX2 and FEN1 were increased in macrophages residing in the infarct, and silencing these nucleases suppressed inflammation, which is critical for post-MI pathogenesis, we selectively silenced *Apex2* and *Fen1* in macrophages after MI using a lipidoid nanoparticle approach (**Figure S6A-6B**). Ejection fraction was improved after the treatment compared to *siControl* at 28 days post-MI (**Figure 3F**). However, systolic volume and cardiac output in the *siApex2* and *siFen1* groups were unchanged compared to the control group (**Figure S6C**). The *siApex2* and *siFen1* groups also exhibited lesser cardiomyocyte hypertrophy (**Figure 3G**). However, the treatment did not significantly suppress collagen deposition in the infarct (**Figure S6D**).

### Myeloid-STING deficiency worsens cardiac function after MI

STING is a pattern recognition receptor, activated by cyclic dinucleotides (cGAMP) produced by cyclic GMP – AMP synthase (cGAS) ^3,6,8^. The cGAS-STING pathway is an arm of the innate immunity that senses nucleic acids in the cytoplasm ^3,4,8^. To examine how mitochondrial biogenesis increases STING activation, we discerned p-STING levels in *Tfam* and *Cmpk2*-deficient macrophages. While *Cmpk2* deficiency lowered oxLDL-induced augmentation of p-STING levels, interestingly, *Tfam*-deficient macrophages had heightened p-STING (**Figure S6E**), confirming the finding that TFAM is crucial for mitochondrial DNA compaction ^38,39^.

Upon activation, STING recruits tank-binding kinase 1 (TBK1), which phosphorylates STING and transcription factor IRF3 to activate the transcription of type I interferon genes ^8,40,41^. However, phospho-TBK1 and phospho-IRF3 levels were comparable between sham and MI cardiac macrophages (**Figure S7A-S7B**). Next, we examined STING activation in cardiac macrophages in the hearts of mice and humans with MI. Macrophage p-STING expression was significantly greater in human (**Figure 4A**) and mouse (**Figure 4B**) hearts with MI compared to the controls. To understand the role of myeloid STING in cardiac remodeling after MI, we generated myeloid-specific *Sting*-deficient mice. Bone marrow-derived macrophages from *LyzM^Cre^*^/+^ *Sting^fl^*^/fl^ animals exhibited lesser inflammatory gene expression compared to the ones isolated from *LyzM*^+/+^ *Sting^fl^*^/fl^ mice (**Figure S7C**). Based on these results, and since inflammation facilitates post-MI fibrosis and ventricular dilation, we hypothesized that the deletion of myeloid *Sting* would be cardioprotective after MI. Interestingly, in contrary to our hypothesis, *LyzM^Cre^*^/+^ *Sting^fl^*^/fl^ animals exhibited lesser left ventricular ejection fraction, global longitudinal strain and a trend towards exacerbated ventricular dilation inferred from systolic and diastolic volumes (**Figure 4C**). Stroke volume and cardiac output were comparable between *LyzM^Cre^*^/+^ *Sting^fl^*^/fl^ and *LyzM*^+/+^ *Sting^fl^*^/fl^ animals. However, *LyzM^Cre^*^/+^ *Sting^fl^*^/fl^ and *LyzM*^+/+^ *Sting^fl^*^/fl^ animals had comparable heart function at baseline (**Figure S7D**). Cardiomyocyte length, diameter, and area were higher in *LyzM^Cre^*^/+^ *Sting^fl^*^/fl^ mice, compared to *LyzM*^+/+^ *Sting^fl^*^/fl^ mice (**Figure 4D-4E**), supporting exacerbated LV dilation. Cardiac fibrosis was significantly expanded in *LyzM^Cre^*^/+^ *Sting^fl^*^/fl^ hearts, compared to *LyzM*^+/+^ *Sting^fl^*^/fl^ animals assessed by Masson’s Trichrome staining (**Figure 4F**) and immunostaining for Collagen1 (**Figure 4G**). The number of TUNEL^+^ apoptotic cells was greater in the infarct of *LyzM^Cre^*^/+^ *Sting^fl^*^/fl^ mice compared to that in *LyzM*^+/+^ *Sting^fl^*^/*fl*^ mice (**Figure S8A**). Finally, we assayed inflammatory cytokine levels in *LyzM^Cre^*^/+^ *Sting^fl^*^/*fl*^ and *LyzM*^+/+^ *Sting^fl^*^/f*l*^ mice after MI. Contrary to reduced inflammatory gene expression *in vitro*, we observed augmented levels of IL-17, IL12-p70, and TNF-⍺ in the serum of *LyzM^Cre^*^/+^ *Sting^fl^*^/fl^ mice after MI, compared to *LyzM*^+/+^ *Sting^fl^*^/fl^ controls (**Figure 4H**). However, the abundance of the immune cell populations in the heart, blood, bone marrow, and spleen was similar between *LyzM^Cre^*^/+^ *Sting^fl^*^/fl^ and *LyzM*^+/+^ *Sting^fl^*^/fl^ mice after MI (**Figure S8B-E**).

**Figure 4:**
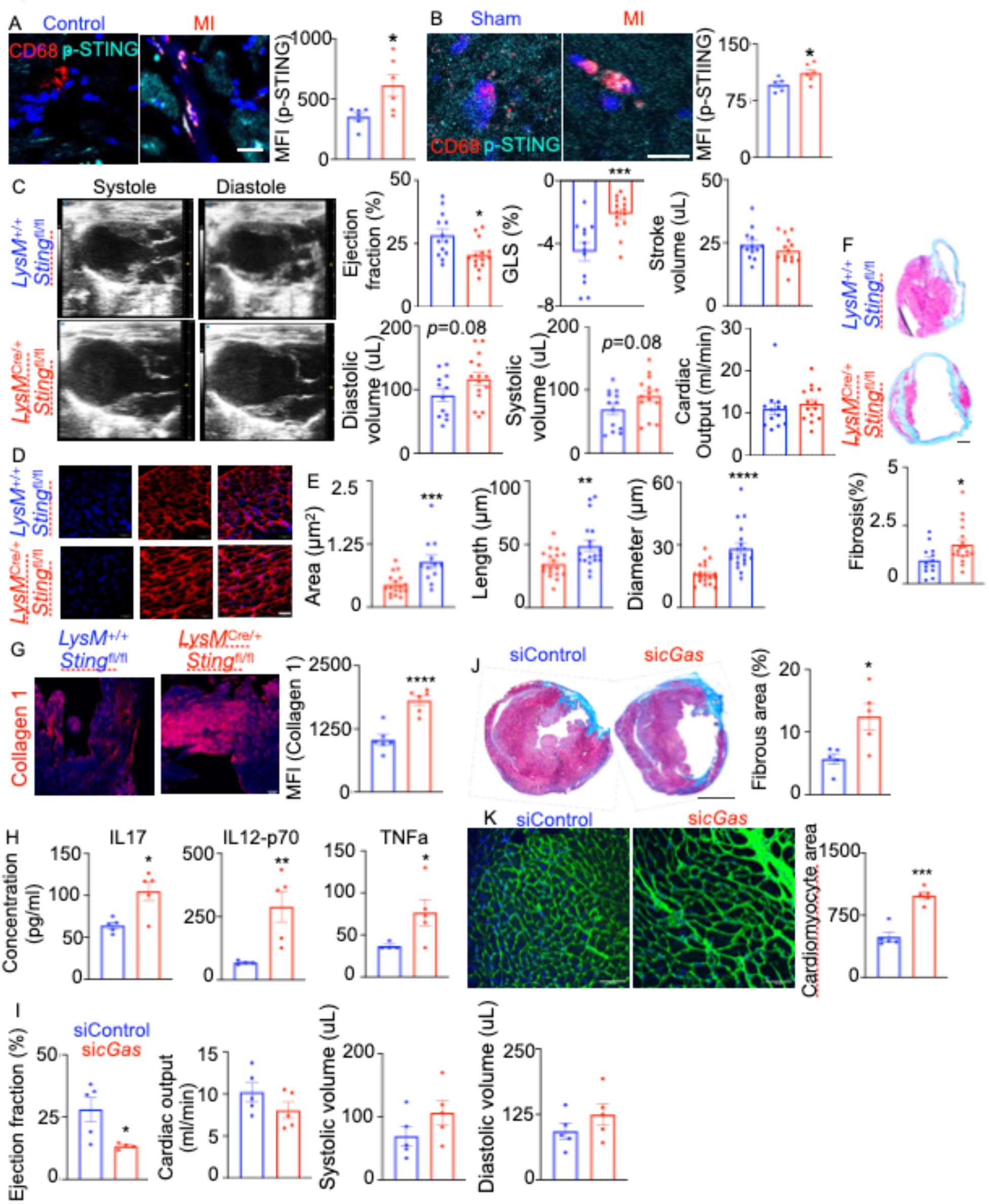
Myeloid *Sting*-deficient mice have exacerbated cardiac remodeling after MI. A-B) Confocal microscopy was conducted to ascertain pSTING expression in cardiac macrophages of control and MI patients (scale bar: 25 μm, n=6/group) (A) and mice with sham and MI surgeries (B, scale bar: 12.5μm, n=5-6/group) (B). C) Echocardiography was performed in *LyzM*^+/+^ *Sting*^fl/fl^ and *LyzMCre*^/+^ *Sting*^fl/fl^ mice on day 28 after MI to evaluate cardiac function (n=12 in *LyzM*^+/+^ *Sting*^fl/fl^ and 14 in *LyzMCre*^/+^ *Sting*^fl/fl^). D-E) Cardiomyocyte area, length, and diameter were calculated by wheat germ agglutinin staining on day 28 after MI (n=15-19/group, scale bar:50μm). F-G) Fibrosis using Masson trichrome staining (F) and Collagen 1 deposition (G) were assessed in the hearts on day 28 after MI (F, scale bar: 500 μm, n=13-18/group; G, scale bar:100 μm, n=6/group). H) A Luminex assay was performed to measure the inflammatory cytokine levels in the serum of and *LyzM*^+/+^ *Sting*^fl/fl^ and *LyzMCre*^/+^ *Sting*^fl/fl^ mice on day 28 after MI (n=5/group). I-K) C57BL/6 mice were injected with either siControl or si*cGas* incorporated in lipidoid nanoparticles for 28 days after MI. Cardiac function by echocardiography (I, n=5/group), cardiac fibrosis by Masson’s trichrome staining (J, scale bar: 500 μm, n=5/group), and cardiomyocyte size by wheat germ agglutinin staining (K, scale bar: 50 μm, n=5/group) were evaluated. \**p*<0.05, \*\**p*<0.01, \*\*\**p*<0.001, \*\*\*\**p*<0.0001, two-tailed Student’s t-test.

Detection of foreign DNA by STING is mediated by the cytoplasmic DNA sensor cGAS, which binds to double stranded DNA in cytoplasm and catalyzes the production of 2’3’ cyclic GMP-AMP (2’3’ cGAMP). This second messenger binds and activates STING ^3,42^. To understand the function of cGAS in macrophages following MI, we silenced *cGas* in macrophages using lipidoid nanoparticles (Figure S9A). *cGas* silencing in macrophages resulted in lower expression of *Sting* and the interferon-responsive genes *Ifih1* and *Ifi27* in macrophages but not in monocytes (**Figure S9B-S9C**). Similar to *LyzM^Cre^*^/+^ *Sting^fl^*^/fl^ mice with MI, *sicGas*-injected mice had depressed ejection fraction, a trend towards greater ventricular dilation (**Figure 4I**), augmented cardiac fibrosis (**Figure 4J**), and expanded cardiomyocyte size (**Figure 4K**).

### Single-cell RNA sequencing discovered STING-regulated genes and pathways in cardiac macrophages after MI

To delineate the mechanisms of cardioprotective function of macrophage STING, we performed single-cell RNA sequencing on the hearts from *LyzM*^+/+^ *Sting^fl^*^/fl^ and *LyzM^Cre^*^/+^ *Sting^fl^*^/fl^ mice after sham surgery and MI (**Figure 5A**). Comparison of the transcriptomic profiles of cardiac macrophages showed upregulation of the genes *Ctsb*, *Stab1*, and *App*, and the transcription factors *Maf* and *Mafb* in *LyzM^Cre^*^/+^ *Sting^fl^*^/fl^ mice after MI (**Figure S10**). Our analyses identified 36 genes common across two comparisons: *LyzM*^+/+^ *Sting^fl^*^/fl^ mice MI versus sham-operated controls, and *LyzM*^+/+^ *Sting^fl^*^/fl^ vs. *LyzM^Cre^*^/+^ *Sting^fl^*^/fl^ after MI (**Figures 5B-5C**). These genes, such as *App, Maf, Nfkb1, Stab1, Ctsb, CD74,* and *Dab2*, are differentially expressed by cardiac macrophages after MI and controlled by STING. Pathway analyses of these differentially expressed genes revealed several biological pathways crucial for cardiac remodeling after MI, such as IL-10 signaling, macrophage alternative activation signaling, dilated cardiomyopathy, collagen degradation, assembly of collagen fibers, and wound healing signaling (**Figure 5D**). Interestingly, the glycation signaling pathway was one of the most significantly altered pathways (Z score= -2). The genes in this pathway, such as *App, Maf, Nfkb,* and *Stab1*, which are differentially expressed in *Sting*^-/-^cardiac macrophages after MI (**Figure 5C**), can lead to cellular responses like inflammation, apoptosis, and tissue remodeling ^43–45^, which determine MI pathogenesis.

**Figure 5:**
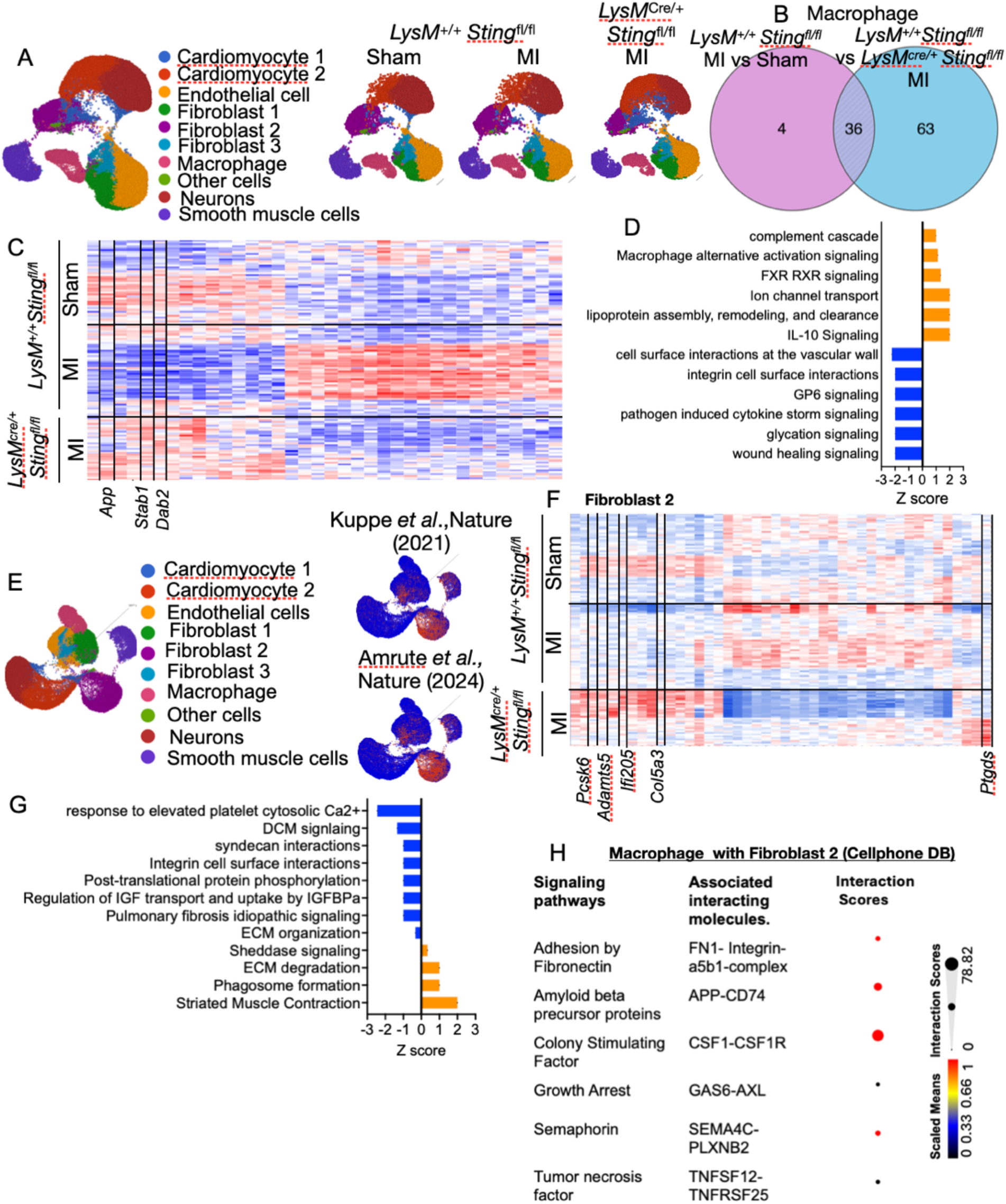
A single-cell transcriptomics study unveils *Stin*g-mediated *App* suppression in cardiac macrophages. A single cell transcriptomics analysis was performed in the hearts of *LyzM*^+/+^ *Sting*^fl/fl^ mice with sham and MI surgeries and *LyzM*^Cre/+^ *Sting*^fl/fl^ mice with MI (n=4/group). A) The UMAP plots represent single-cell transcriptomic data showing various cell types in the hearts defined by transcriptomic signatures. B) The Venn diagram displays the STING-controlled genes, which are differentially expressed in macrophages after MI. C) The heatmap depicts these 36 genes. D) Ingenuity Pathway Analyses (IPA) of these genes were carried out. E) Pathological fibroblast populations in the hearts after MI were identified by analyzing the myofibroblast genes identified by Amrute et al. and Kuppe et al. F) The heatmap displays the genes differentially expressed in Fibroblast 2 in various groups. G) A pathway analysis of the genes differentially expressed in Fibroblast 2 was conducted. H) Macrophage–Fibroblast 2 cell communication analyses by Cellphone DB show the signaling pathways activated in the absence of myeloid Sting. n=4/group.

Because our data unveiled exaggerated fibrosis in the heart in mice lacking myeloid *Sting* after MI, we sought to decipher the changes in the transcriptomic signatures of cardiac fibroblast in *LyzM^Cre^*^/+^ *Sting*^fl/fl^ mice with MI. Three distinct fibroblast populations were identified by our analyses (**Figures 5A, S11A, and S11B**). To discern pathogenic fibroblast(s) after MI, we compared our dataset to published studies ^46,47^. Fibroblast 2 was enriched with the genes expressed by pathogenic fibroblasts reported by Kuppe et al. and FAP and POSTN-expressing Fibroblast 9 reported by Amrute et al. ^46,47^ (**Figure 5E**). Sting-deficiency in myeloid cells resulted in significantly altered expression of *Col5a3, Ifi205, Ptgds, Pcsk6, and Adamts5* in Fibroblast 2 (**Figure 5F**). These genes are crucial in matrix remodeling and organization, inflammation and immune cell activation. Fibroblast 2 in *LyzM^Cre^*^/+^ *Sting*^fl/fl^ mice with MI displayed many suppressed biological pathways, including fibrosis, protein phosphorylation, dilated cardiomyopathy signaling, and extracellular matrix organization, which aggravated ventricular dilation after MI (**Figure 5G**). Given the enrichment of pathogenic fibroblast genes in Fibroblast 2, we explored the interactions of macrophage and Fibroblast 2 using Cellphone DB (**Figure S12**). We observed that APP-CD74 and CSF1-CSF1R had high interaction scores between these two cell populations (**Figures 5H**).

### Myeloid STING regulates App to limit remodeling and preserve heart function after MI

Because amyloid precursor protein (APP) expression was upregulated in *Sting*^-/-^ cardiac macrophages after MI (**Figures 5, 6A, and 6B**), and APP-CD74 had a significant interaction score between cardiac macrophage and Fibroblast 2, we sought to probe the function of APP in cardioprotection mediated by myeloid STING. To rule out developmental effects of myeloid STING and target this gene specifically in macrophages but not neutrophils, we employed lipidoid nanoparticles to deliver siRNA against *Sting* or both *Sting* and *App* (**Figure S13**). Lipidoid nanoparticles-assisted siRNA delivery mediated gene silencing in macrophages but not in neutrophils, monocytes, and B cells (**Figure S13A-S13F**). As expected, *siSting* treatment significantly decreased ejection fraction, stroke volume, and cardiac output at 4 and 8 weeks post-MI, in concordance with our observation in *LyzM^Cre^*^/+^ *Sting*^fl/fl^ mice after MI (**Figure 6C**). However, *App* silencing abolished the detrimental effects on heart function due to the silencing of macrophage *Sting*. Congruently, *App* silencing in macrophages reversed exuberant cardiac fibrosis instigated by *Sting* silencing in macrophages after MI (**Figure 6D**). Leukocyte frequencies in blood, spleen, and heart were examined by flow cytometry. The s*iSting* and *siSting+siApp* groups exhibited a reduction in blood and spleen monocytes, while there was an increase of neutrophils in the heart of the *siSting+siApp* group (**Figure S14 A-C**). To test how APP induces fibrosis, we cultured fibroblasts in presence of the supernatants obtained from *Sting*^-/-^ BMDM after transfecting either siControl or si*App*. As expected, the supernatant of *Sting*^-/-^ BMDM contained a profuse amount of APP whereas *App* silencing reduced the amount of this protein in the supernatant (**Figure 6E**). *Sting* deficiency in macrophages drove fibroblasts into the cell cycle and increased the expression of the genes regulating fibrosis (**Figures 6F, 6G, and S14D**). In contrast, *App* silencing in BMDM reversed these responses in fibroblasts (**Figure 6F, 6G, and S14D**).

**Figure 6:**
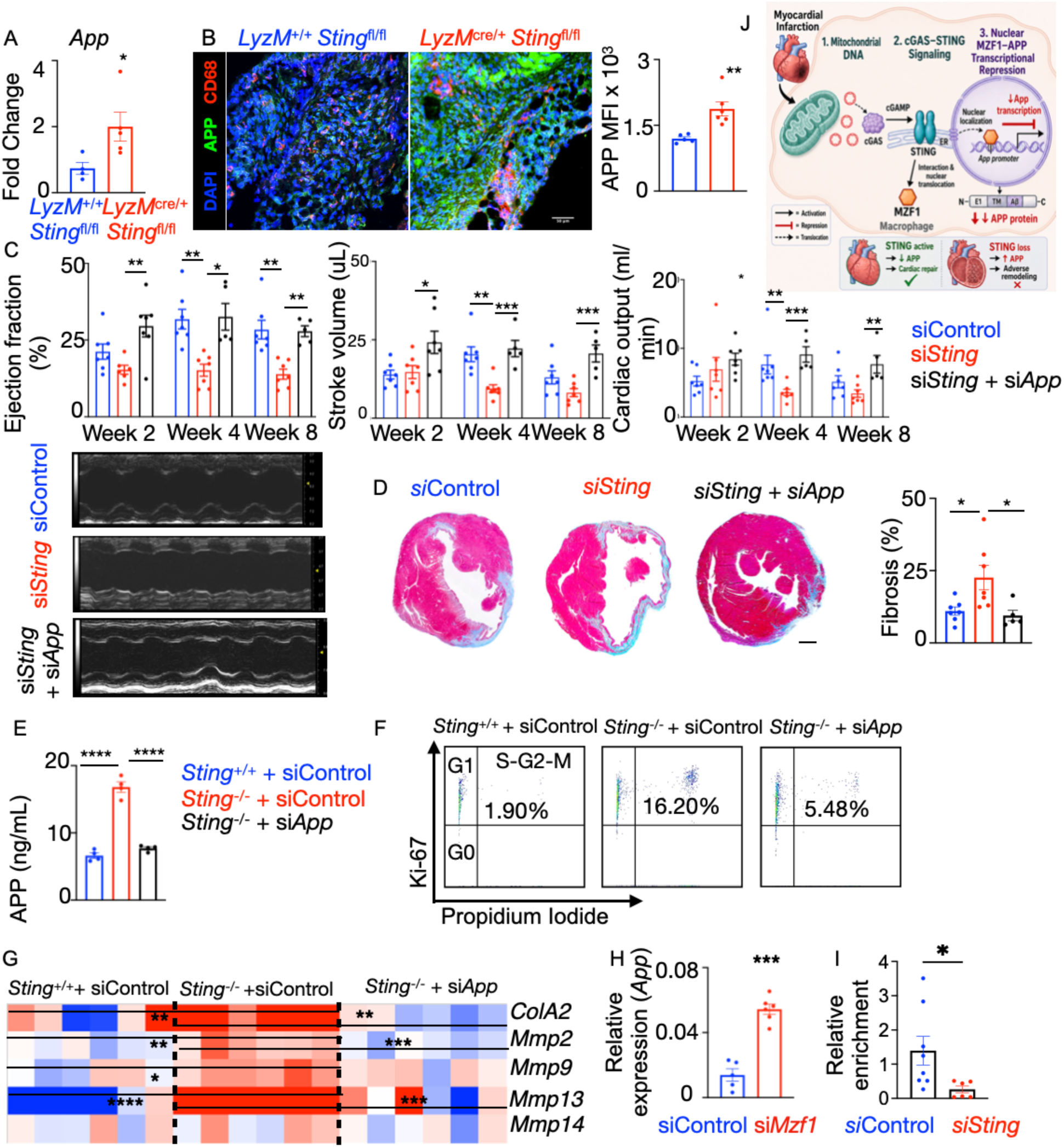
Myeloid STING suppresses APP to mediate cardioprotective effect after MI. A-B) APP was quantified in *Sting*^+/+^ and *Sting*^-/-^ BMDM by qPCR (A, n=4/group) and cardiac macrophages after MI in *LyzM*^+/+^ *Sting*^fl/fl^ and *LyzM^Cre^*^/+^ *Sting*^fl/fl^ mice after MI (B, scale bar: 50μm, n=6/group). C-D) C57BL/6 mice were treated with either siControl, si*Sting*, or si*Sting*+si*App* incorporated in lipidoid nanoparticles after MI. Cardiac function by echocardiography at 2, 4, and 8 weeks after MI (C, n=7/group) and cardiac fibrosis by Masson’s trichrome staining (D, scale bar: 500 μm) were evaluated. E) BMDM were treated with either *siControl*, *siSting* or *siSting+siApp*, and APP was measured in the supernatant (n=4/group). F-G) 3T3-L1 fibroblasts were cultured with conditioned media collected from these macrophages. Cell proliferation by flow cytometry (F, n=6 per group) and pro-fibrotic gene expression by qPCR (G, n=6/group) were ascertained. H) qPCR was performed to quantify *App* in BMDM after *Mzf1* silencing. n=5/group. I) ChIP was performed to assess the enrichment of MZF1 on the *App* promoter after *Sting* silencing in BMDM. n=6-8/group. \**p*<0.05, \*\**p*<0.01, \*\*\**p*<0.001, \*\*\*\**p*<0.0001, two-tailed Student’s t-test (A, B, H, and I), one-way ANOVA with post-hoc Fisher LSD test (C-G).

To examine the molecular mechanisms of the control of *App* expression by STING, we performed an *in silico* analysis to identify the transcription factors (TFs) that bind to the *App* promoter. Our analysis revealed MZF1, NFAT, ING4, YY1, KLF4, ZNF333, AhR, and TTF-1 as possible TFs interacting with the *App* promoter. Among these TFs, MZF1, ZNF333, and TTF-1 are known suppressors. We observed that *Mzf1* and *Nfatc1* silencing in BMDM significantly increased *App* expression while Ahr silencing decreased *App* (**Figure 6H and S14E**). A proximity ligation assay demonstrated STING-MZF1 interaction, and *Sting* deficiency significantly reduced MZF1 nuclear colocalization (**Figure S15A**). Additionally, *Sting*^-/-^ BMDM had reduced MZF1 levels (**Figure S15B**). Finally, using ChIP-qPCR, we experimentally validated MZF1 binding motif on the *App* promoter and observed that the enrichment of MZF1 on the *App* promoter is reduced upon *Sting* silencing (**Figure 6I**). In summary, these findings suggest that STING interacts with MZF1 and facilitates its nuclear translocation, resulting in MZF1-mediated suppression of *App* transcription (**Figure 6J**).

## Discussion

Ischemic heart disease caused by myocardial infarction (MI) is among the leading causes of morbidity and mortality across the world ^48^. Inflammatory pathways play a major role in healing, remodeling, and the progression of heart failure after MI ^49,50^. Macrophages comprise one of the largest immune cell populations in the heart and play a wide range of roles from cardiac homeostasis to chronic inflammation and adverse remodeling in heart failure (HF) ^51,52^ via secretion of inflammatory cytokines and modulating ECM production ^53–55^. Although STING has been shown to propagate inflammation, our data demonstrate that macrophage STING is beneficial after MI. Specifically, we demonstrate that STING suppresses APP production by translocating MZF1 into the nucleus, suppressing fibrosis and cardiac remodeling.

We also show that uncontrolled synthesis of mitochondrial DNA, which is known to stimulate the STING pathway, in infarct macrophages contributes to exacerbated inflammation and worse outcome after MI. Our data reveal that macrophages exposed to an inflammatory stimulus or cardiac macrophages in ischemic hearts have greater levels of oxidative DNA damage and augmented mtDNA release into the cytoplasm. This is consistent with an expanding body of literature on mitochondrial DNA as a proinflammatory stimulus and the role of mitochondrial dynamics in modulating macrophage function ^2,24,33^. Damaged mtDNA is cleaved by mitochondrial nucleases, such as FEN1 and APEX2, expediting mtDNA release into the cytoplasm and fueling inflammation^56,57^. In line with these findings, our study shows attenuated inflammation and improved cardiac function when these nucleases were silenced in macrophages. It has been shown that mitochondrial DNA, once in the cytosol, stimulates cGAS-STING signaling and promotes NLRP3 inflammasome activation ^35,36^. The ability to detect a broad spectrum of self and foreign nucleic acids makes the cGAS-STING pathway a unique arm of innate immunity and is of therapeutic interest in various autoimmune diseases and cancers.

Danger-associated molecular patterns (DAMPs) secreted after an injury can activate the cGAS-STING pathway and drive inflammation in cardiovascular diseases ^9,58^. Studies have highlighted the mechanistic roles and therapeutic potentials of cGAS-STING signaling in the pathology of cardiovascular diseases ^4,26,58^. Systemic administration of STING inhibitors H-151 or C-178 has been demonstrated to preserve cardiac function after MI and reduce adverse cardiac remodeling and fibrosis ^28,59^. In murine pressure overload models, it has been reported that cGAS inhibition or global deficiency of *Sting* ameliorated inflammation and reduced fibrosis and hypertrophy ^60,61^. The detrimental role of STING as reported in these studies could be driven by its expression by cardiomyocytes. While *Sting*^-/-^ macrophages in vitro exhibited a blunted inflammatory response to oxidized LDL, our study showed that myeloid *Sting*-deficient mice had worse remodeling and cardiac function after MI. In line with our findings, recent studies have shed light on the protective role of myeloid STING. Sulka et al., demonstrated that STING signaling in microglia limits age-associated damaged DNA accumulation and maintains the integrity of the blood brain barrier ^62^. *Sting*^-/-^ microglia from aged mice exhibited higher expression of markers associated with neurodegenerative diseases. Additionally, a recent study showed that STING-YBX1 interaction alleviated thrombus formation in a mouse model of deep vein thrombosis ^63^. In line with the findings of beneficial roles of myeloid STING, our data underscores the significance of myeloid STING in cardiac function after MI. As such, cell-specific STING targeting is required to harness its therapeutic potential.

In addition to their classical role associated with phagocytosis of tissue debris after MI, macrophages secrete chemokines to recruit additional immune cell subsets to the infarct area and promote fibrosis by mediating fibroblast differentiation into myofibroblasts ^64^. While we observed lower expression of inflammatory genes and cytokines in *Sting*^-/-^ macrophages *in vitro* in line with previous reports^27,29,61^, greater levels of inflammatory cytokines were observed in the serum of *LyzM*^Cre/+^ *Sting*^fl/fl^ mice after MI along with exacerbated cardiac fibrosis and left ventricular dilation. Consistent with our unexpected finding of higher degree of inflammation and fibrosis in absence of *Sting in vivo*, a recent publication showed elevated type I interferon response in microglia lacking *Sting* due to accumulation of neuronal DNA damage ^62^. Given the studies reporting deleterious roles of STING in various cell types including cardiomyocytes and endothelial cells^20,24,28,29,59^, the cardioprotective effects of macrophage STING after MI are unexpected and interesting. Similarly, STING expressed by dendritic cells mediates anti-tumor immunity by activating glycolysis^11^, whereas B cell STING elicits pro-tumor effects by suppressing NK cells^31^. Therefore, STING provokes contrasting physiological effects depending on cell type involved. However, we do not know why STING expressed by cardiomyocytes and macrophages exerts opposing functions in the heart. Different epigenetic modifications can lead to diverse functions of a protein depending on cell types^65,66^. Besides, a recent study showed that various cGAS-like receptors mediate diverse STING functions^67^. Future studies are warranted to compare epigenetic modifications of STING and cGAS-like receptors in cardiomyocytes and cardiac macrophages to explain the cell type-specific effects of STING on cardiac remodeling. Additionally, we interestingly found that MI did not alter pTBK1 and pIRF3 in macrophages, indicating that myeloid STING-mediated cardioprotection is independent of these two proteins. This finding is in line with the report of TBK1-independent non-canonical functions of STING ^68^.

To probe the mechanisms of enhanced cardiac fibrosis in myeloid *Sting-*deficient mice, we computationally analyzed cardiac macrophage-fibroblast interaction and observed that APP propels post-MI signaling from *Sting*^-/-^ macrophages to myofibroblasts. Congruently, *in vivo* silencing of *App* in macrophages ameliorated the pathological effect of myeloid *Sting* deficiency in the heart. Amyloid precursor protein APP is a transmembrane protein known to be highly expressed in neural synapses. Although *App* is expressed in a wide variety of cell types including macrophages ^45^, it is well studied in the context of Alzheimer’s disease. Macrophages can secrete soluble APP ^69^, which can affect the functions of other cells in a paracrine manner. The significance of this protein expressed in macrophages is relatively understudied in cardiovascular disease. Interestingly, APP plays an immunoregulatory role in macrophage responses to high fat diet and LPS ^45^. Furthermore, APP-CD74 interaction is crucial for paracrine communication between endothelial cells and macrophages, and promotes fibrotic gene expression ^70^. Downregulation of *APP* dampens the inflammatory response to LPS and secretion of inflammatory cytokines ^69^. Our findings show a novel non-canonical function of STING in cardiac macrophages, highlighting the crosstalk between the nucleic acid sensing pathways and APP. Inflammation-independent roles of STING in nociception ^71^, autophagy ^72^, lysosomal biogenesis ^68,73^, and polyunsaturated fatty acids ^74^ have been reported by recent publications. Although our analysis suggests that the pro-fibrotic effect of APP is mediated via CD74 expressed by cardiac fibroblasts, a Cre recombinase mouse strain lacking CD74 specifically in fibroblasts will be required to test this interesting hypothesis.

## Author contributions

NN, EJ and PD designed the experiments, analyzed the data, and wrote the manuscript. VS, MH, PAA, AD, AR, LOS assisted with the experiments. PD conceived and supervised the project, provided funding for the project, and wrote the manuscript.

## Acknowledgements

This research was funded by the National Institutes of Health (NIH) grants R01HL142629, R01HL142629-04W1, R01HL143967, R01AG069399 and R01DK129339, the VA Merit Award 1I01BX006392-01, the American Heart Association (AHA) Transformational Project Award (19TPA34910142), the AHA Innovative Project Award (19IPLOI34760566), the ALA Innovation Project Award (IA-629694), the AHA Innovative Project Award (23IPA1053549), the AHA Collaborative Sciences Award (24CSA1257253), and the AHA Established Investigator Award (25EIA1415660) to P. Dutta, NHLBI K99/R00 HL157689 to N. Natarajan and AHA 25POST1368717 to E Johny. Partek Flow software, version 10.0.22.1204, licensed by the HSLS University of Pittsburgh, was used to analyze single-cell transcriptomic data. The University of Pittsburgh Center for Research Computing, RRID:SCR_022735 supplied the resources necessary to support this research, in part. Specifically, this work used the HTC cluster, which is supported by NIH award number S10OD028483. BioRender software was used to create the visual abstract and schematic figures. This work benefitted from Cytek Aurora CS Spectral Sorter funded by NIH S10OD032265.

## Supplementary figures

**Figure S1:**
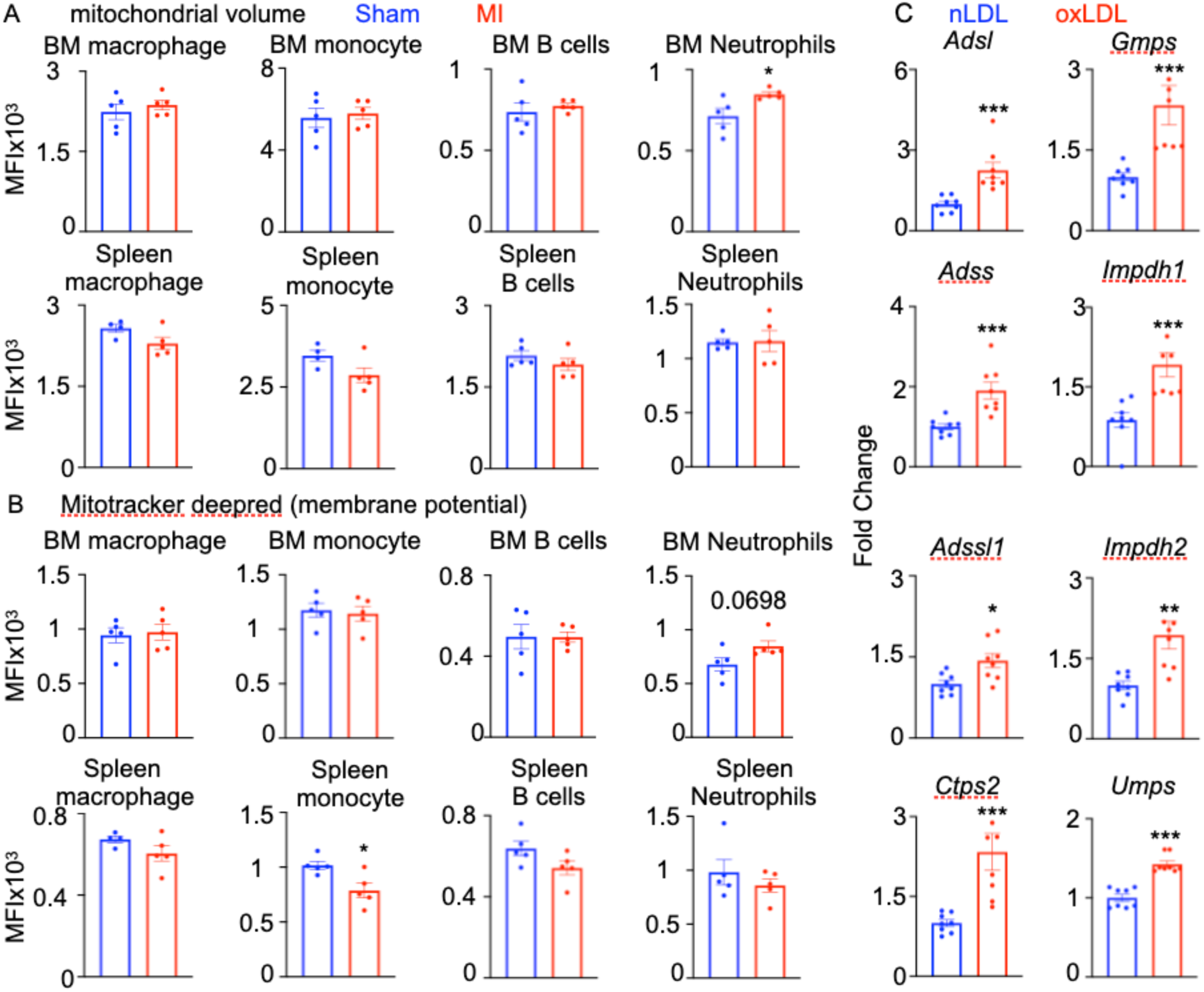
MI does not alter mitochondrial volume and membrane potential in most bone marrow and splenic leukocytes. A-B) Mitochondrial volume using mitotracker green (A) and membrane potential using mitotracker deepred (B) in leukocyte subsets of bone marrow and spleen of sham and MI mice were calculated by flow cytometry, n=5/group. C) Nucleotide biosynthesis genes in oxLDL-treated BMDM were quantified by qPCR, n=7 nLDL, n=8 oxLDL. \**p*<0.05, \*\**p*<0.01, \*\*\**p*<0.001, \*\*\*\**p*<0.0001, two-tailed Student’s t-test.

**Figure S2:**
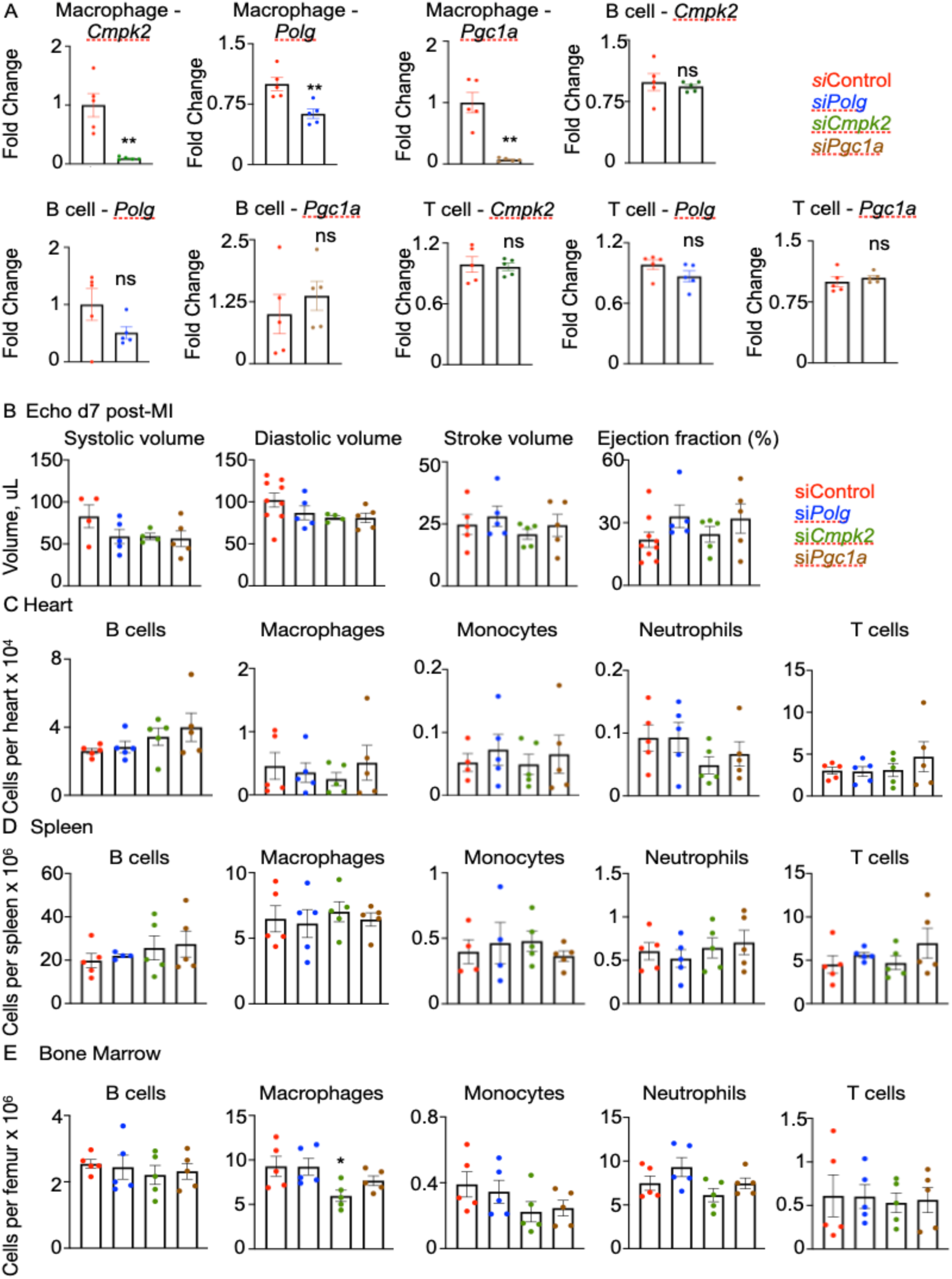
Silencing of mitochondrial biogenesis genes in macrophages after MI improves cardiac function. *siControl, siPolg, siPgc1a,* and *siCmpk2* incorporated in lipidoid nanoparticles were injected i.v in mice after MI. A) Gene expression was quantified using qPCR in the sorted cells to validate macrophage-specific *in vivo* gene silencing. B) Heart function was measured on day 7 after MI using echocardiography. C-E) The leukocyte subsets in the heart (C), spleen (D), and bone marrow (E) were enumerated by flow cytometry on day 28 after MI. n=4-5/ group. \**p*<0.05, \*\**p*<0.01, \*\*\**p*<0.001, \*\*\*\**p*<0.0001, one-way ANOVA with post-hoc Fisher LSD test.

**Figure S3:**
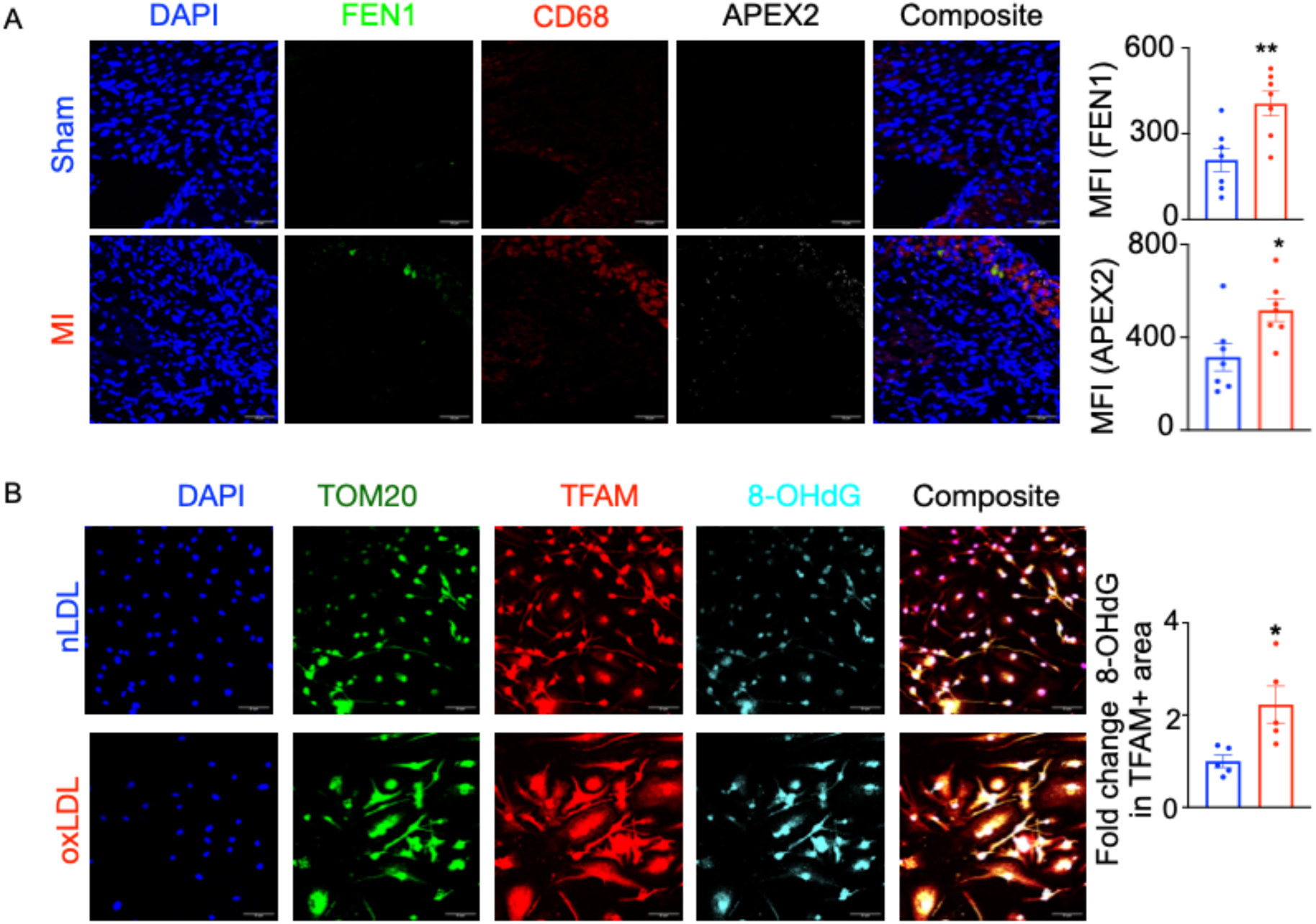
MI heightens mitochondrial nuclease levels in cardiac macrophages. A) Mitochondrial nuclease expression in cardiac macrophages was quantified by immunofluorescence on day 28 after MI, n=6/group. B) oxLDL-induced oxidative damage of mitochondrial DNA in BMDM was assessed by TFAM, 8-OHdG, and TOM20 staining, n=5/group. \**p*<0.05, \*\**p*<0.01, two-tailed Student’s t-test.

**Figure S4:**
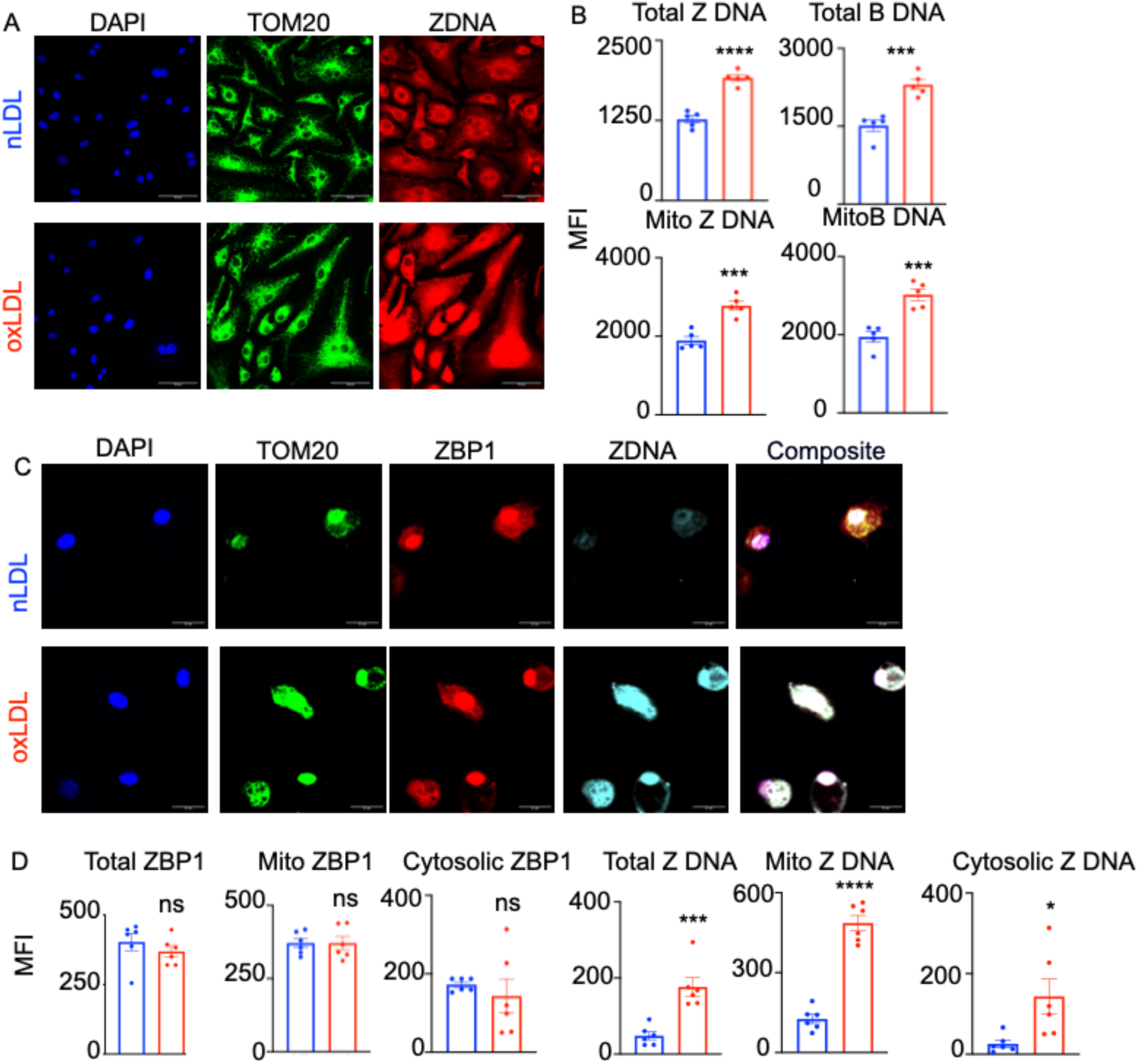
oxLDL treatment augments mitochondrial Z DNA content. Total and mitochondrial B DNA and Z DNA in mouse BMDM (A, B) and human monocyte-derived macrophages (C, D) treated with nLDL or oxLDL were assessed. Scale bar: 50 μm (A), 25 μm (C), n=5/group. \**p*<0.05, \*\**p*<0.01, \*\*\**p*<0.001, \*\*\*\**p*<0.0001, two-tailed Student’s t-test.

**Figure S5:**
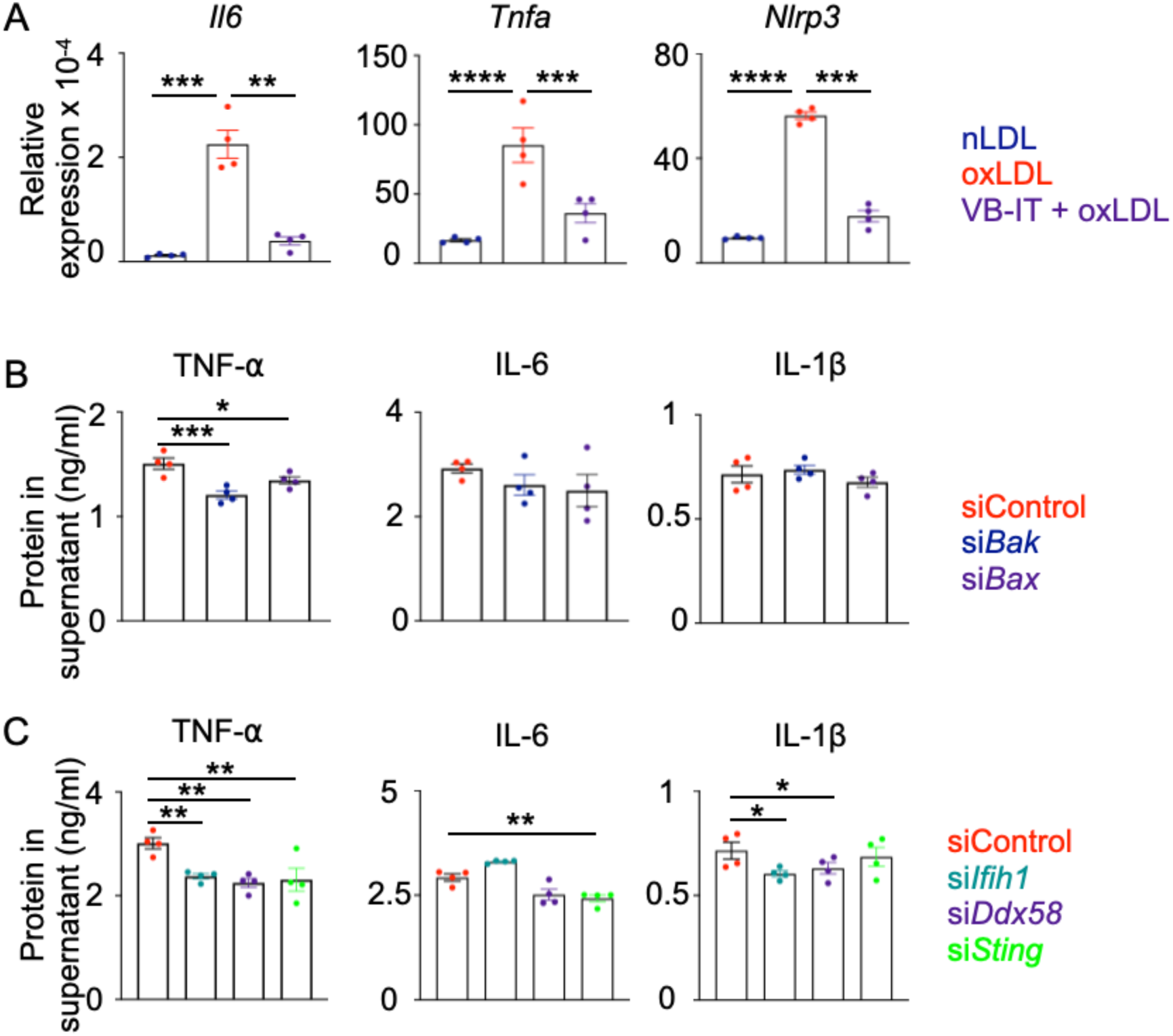
The blockade of mitochondrial DNA exit into the cytoplasm suppresses inflammation. A) Inflammatory response in response to oxLDL in BMDM was examined after VDAC inhibition with VB-IT, n=4/group. B-C) The inflammatory cytokines were quantified using ELISA in BMDM supernatant after silencing *Bak* or *Bax* (B) and the cytoplasmic nucleic acid sensors *Ifih1*, *Ddx58*, and *Sting* (C), n=4/group. \**p*<0.05, \*\**p*<0.01, \*\*\**p*<0.001, \*\*\*\**p*<0.0001, one-way ANOVA with post-hoc Fisher LSD test.

**Figure S6:**
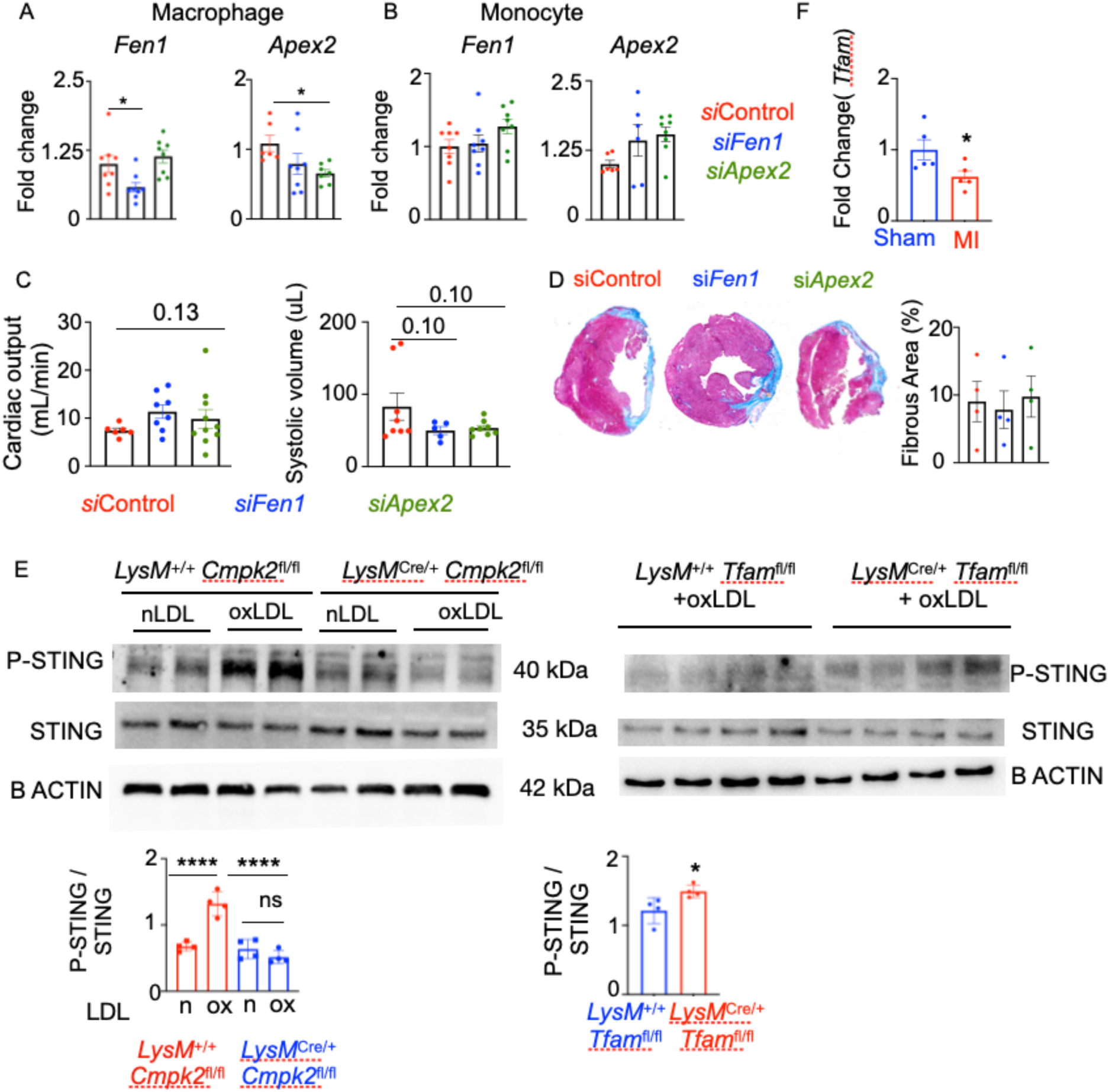
*Cmpk2* deficiency diminishes pSTING expression in BMDM. A-C) siControl, si*Fen1*, or si*Apex2* formulated in lipidoid nanoparticles were injected i.v in C57BL/6 mice after MI. *Fen1* and *Apex2* were quantified in cardiac macrophages (A) and monocytes (B) isolated from mice treated with siRNA against the genes, n=8/group. (C) Echocardiography was performed to assess systolic volume and cardiac output on day 28 after MI, n=6-8/group. D) Fibrosis was examined using Masson’s trichrome staining, Scale: 500 μm, n=4/group.(pSTING levels in *Cmpk2*^-/-^, *Tfam*^-/-^, and wildtype macrophages exposed to either native or oxidized LDL were estimated by immunoblot, n=4/group. \**p*<0.05, \*\*\*\**p*<0.0001, one-way ANOVA with post-hoc Fisher LSD test.

**Figure S7:**
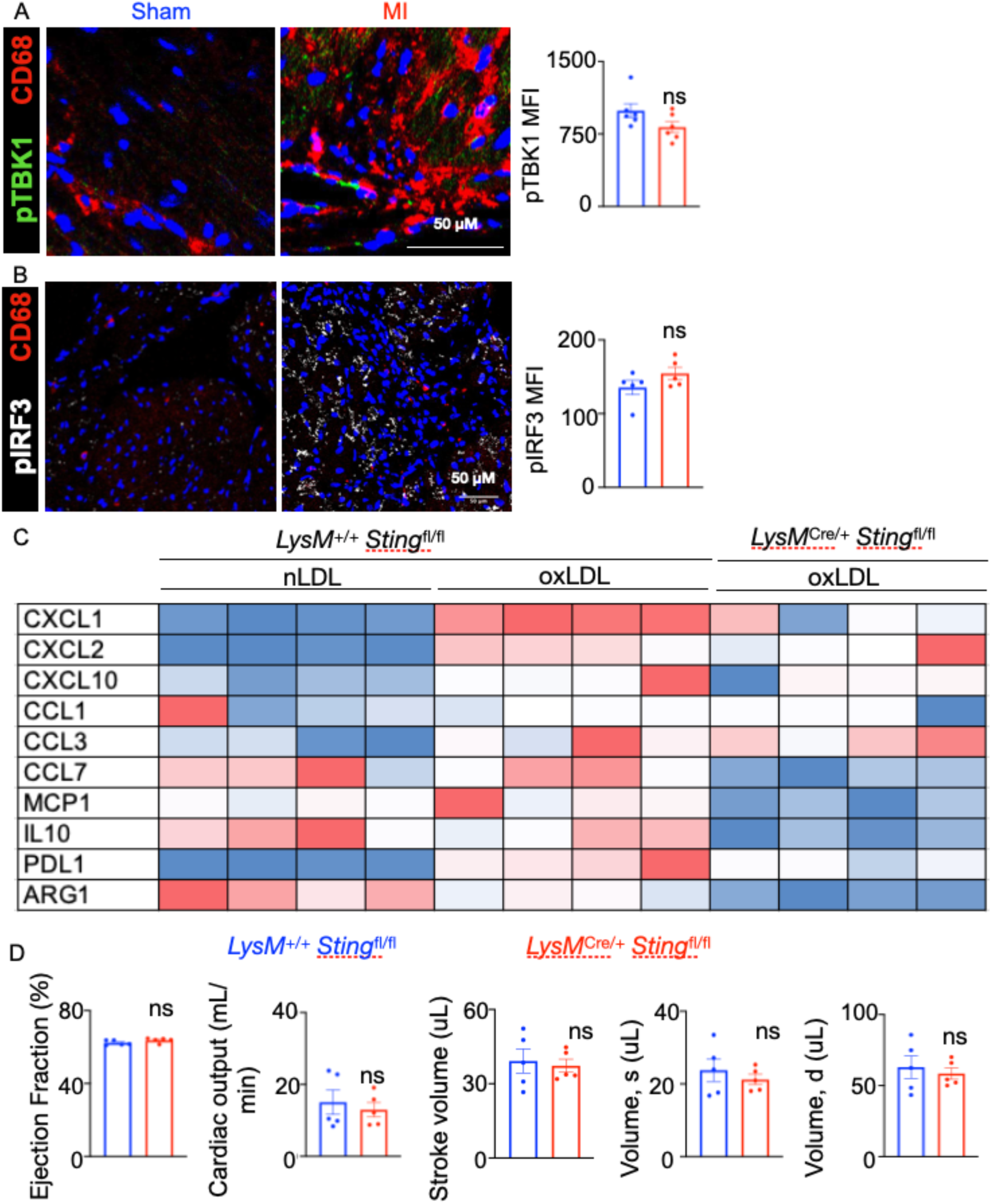
Macrophage *Sting* deficiency suppresses inflammation *in vitro*. A-B) pTBK1 (A) and pIRF3 (B) in cardiac macrophages of mice with sham and MI surgeries were quantified by confocal microscopy. The representative images from 2 experiments are shown on the left and the quantification of the proteins is depicted on the right, n=6 (pTBK1), 5 (pIRF3). C) Expression of inflammatory cytokines and interleukins in *Sting*^+/+^ and *Sting*^-/-^ macrophages treated with nLDL or oxLDL was examined by qPCR, n=4/group. D) Steady state cardiac functional parameters were assessed by echocardiography in *LyzM*^+/+^ *Sting*^fl/fl^ and *LyzM^Cre^*^/+^ *Sting*^fl/fl^ mice, n=5/group, 2-tailed Student’s t-test (A,B,D), one-way ANOVA with post-hoc Fisher LSD test (C).

**Figure S8:**
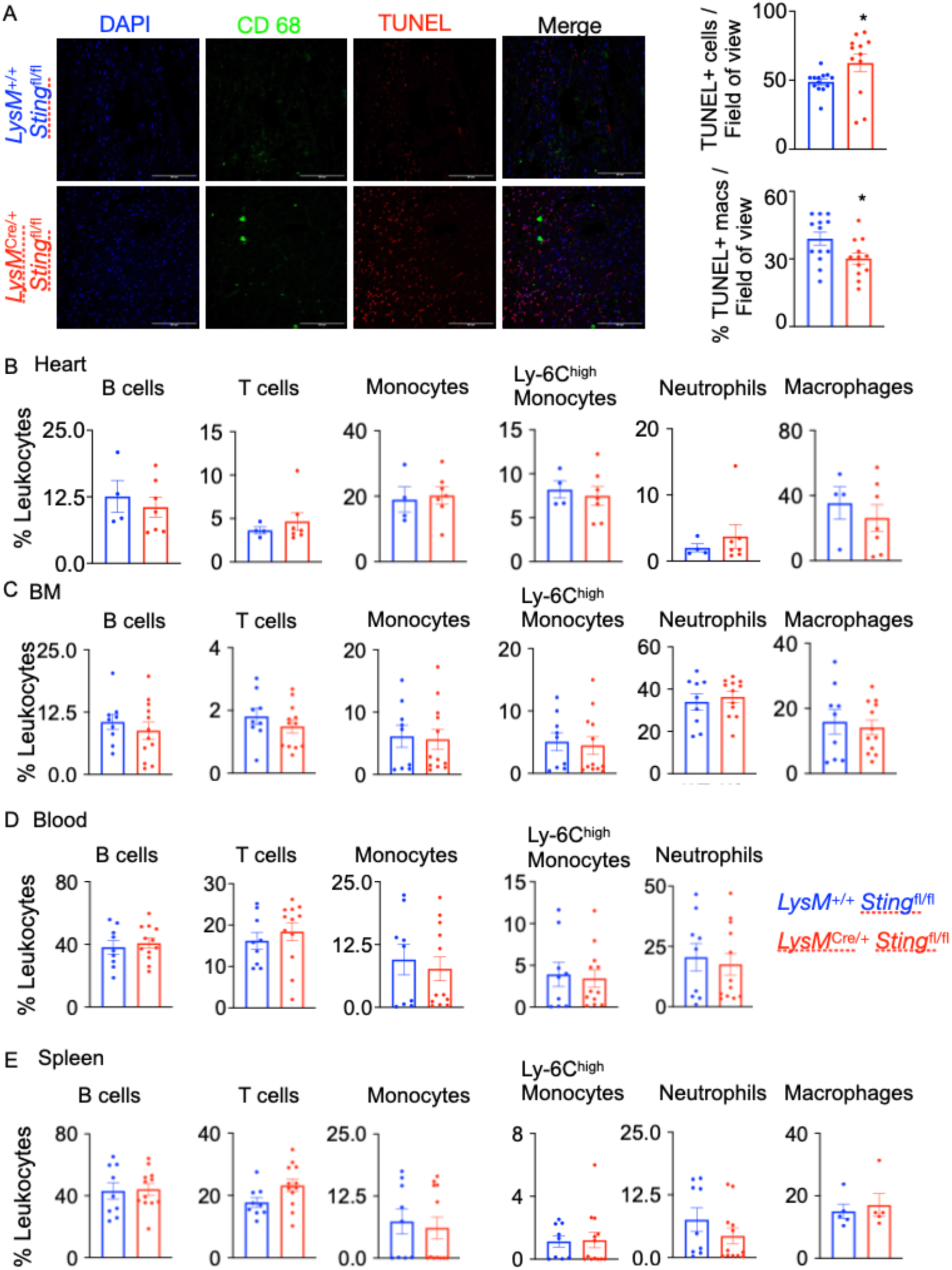
Myeloid *Sting* deficiency does not alter the abundance of leukocyte populations after MI. A) Apoptotic cells in the hearts of *LyzM*^+/+^ *Sting*^fl/fl^ and *LyzM^Cre^*^/+^ *Sting*^fl/fl^ mice after MI were assessed by TUNEL staining, representative images from 2 independent experiments, n=12-13/group. B-E) Leukocyte populations in heart (B), bone marrow (C), blood (D), and spleen (E) of *LyzM*^+/+^ *Sting*^fl/fl^ and *LyzM^Cre^*^/+^ *Sting*^fl/fl^ mice with MI were examined by flow cytometry, n=9-12/group. \**p*<0.05, \*\**p*<0.01, \*\*\**p*<0.001, \*\*\*\**p*<0.0001, 2-tailed Student’s t-test.

**Figure S9:**
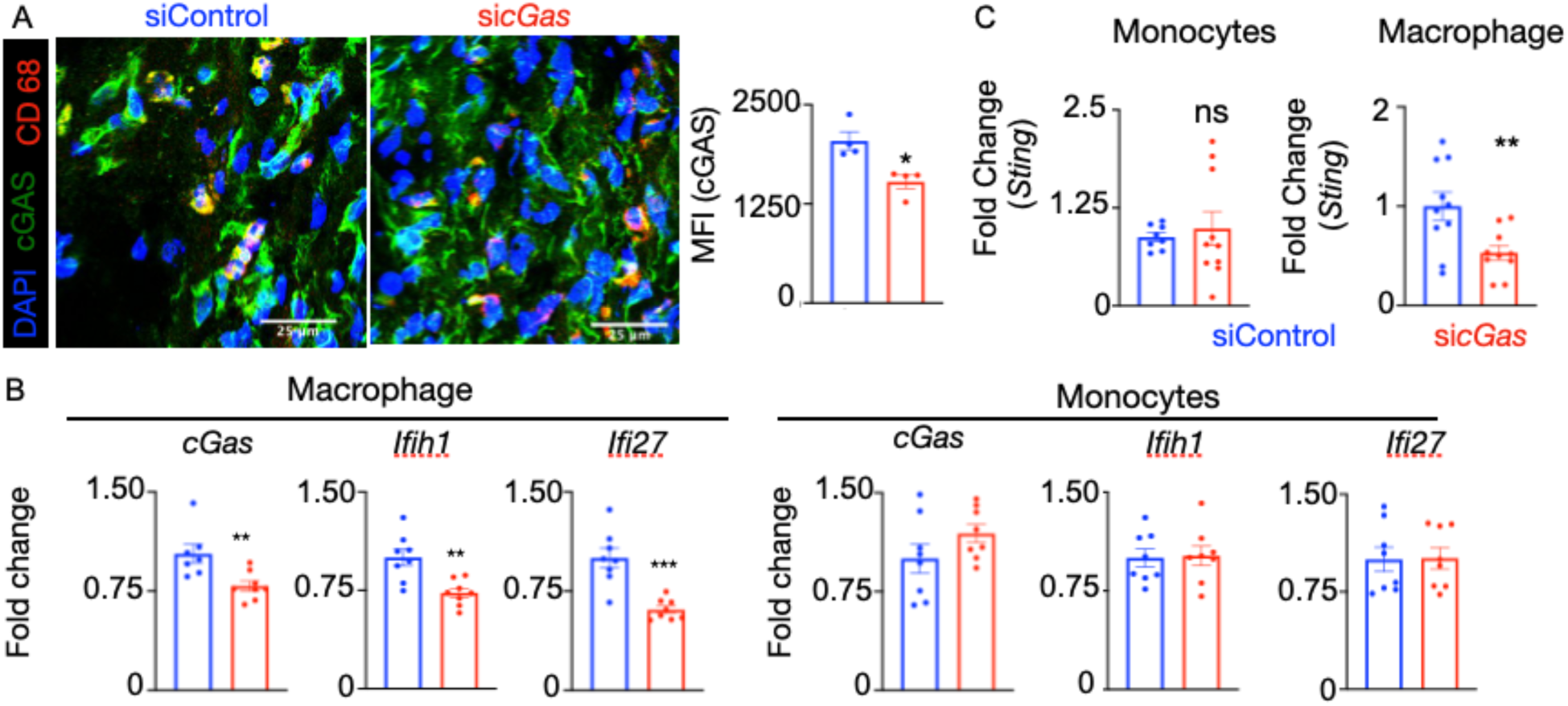
siRNA encapsulated in lipidoid nanoparticles specifically target macrophages. siRNA against *cGas* incorporated in lipidoid nanoparticles were injected i.v in C57BL/6 mice after MI. cGas expression was examined in cardiac sections by confocal microscopy (A). The expression of *cGas*, downstream interferon-responsive genes, such as *Ifih1* and *Ifi27* (B) and *Sting* (C) were quantified in macrophages and monocytes after si*cGas* injection. n=10 (A), 8 (B) and 4 (C)/group. \**p*<0.05, \*\**p*<0.01, 2-tailed Student’s t-test.

**Figure S10:**
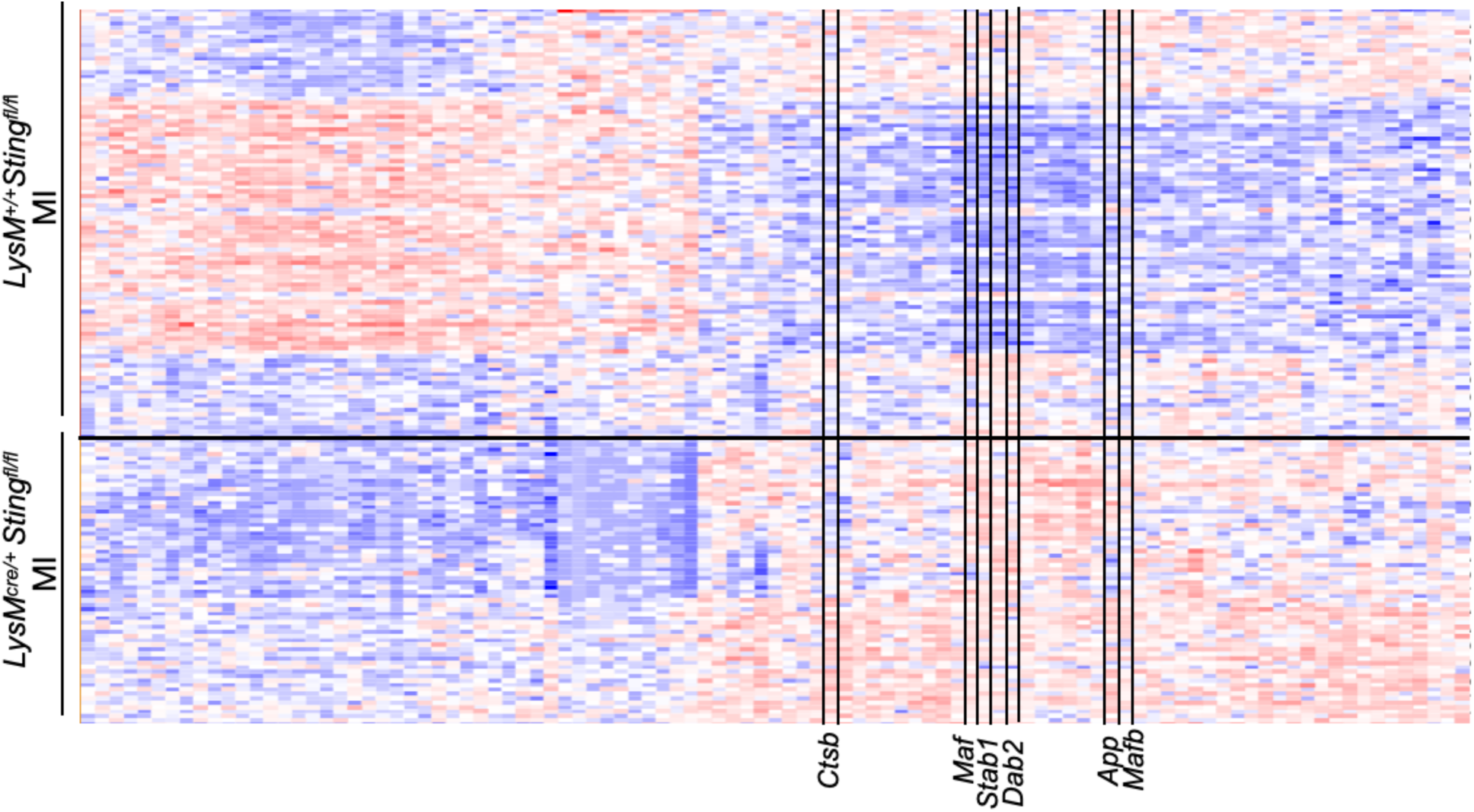
scRNA sequencing reveals transcriptomic differences in *Sting*^-/-^ cardiac macrophages. The heatmap shows the expression of the genes differentially regulated in cardiac macrophages of *LyzM*^+/+^ *Sting*^fl/fl^ and *LyzM^Cre^*^/+^ *Sting*^fl/fl^ mice after MI. n=4/group.

**Figure S11:**
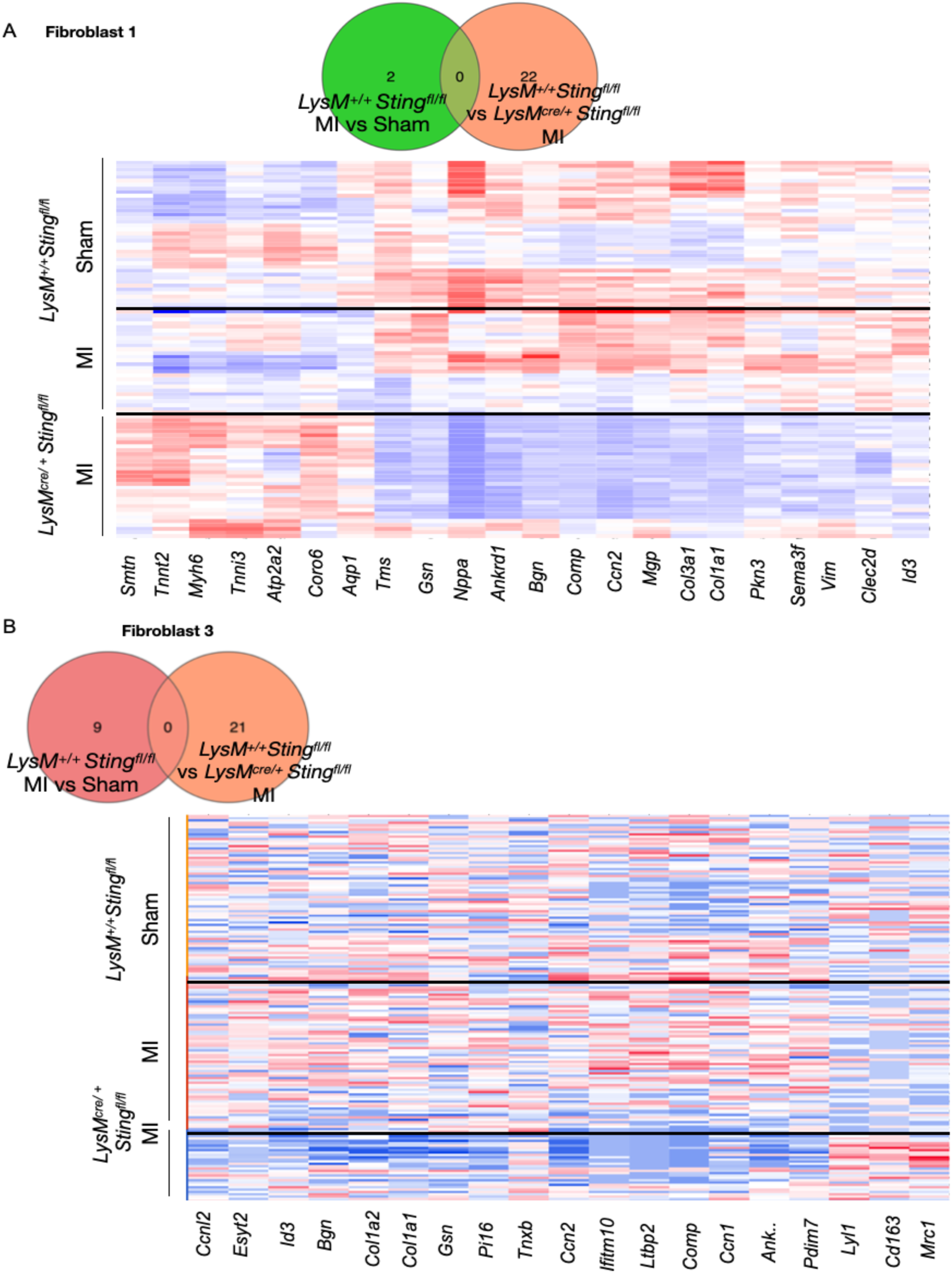
scRNA sequencing reveals transcriptomic differences in cardiac fibroblasts of myeloid-*Sting* deficient mice. Venn diagram and heatmap display the genes differentially expressed in Fibroblast 1 (A) and Fibroblast 3 (B) populations in various groups of mice. n=4/group.

**Figure S12:**
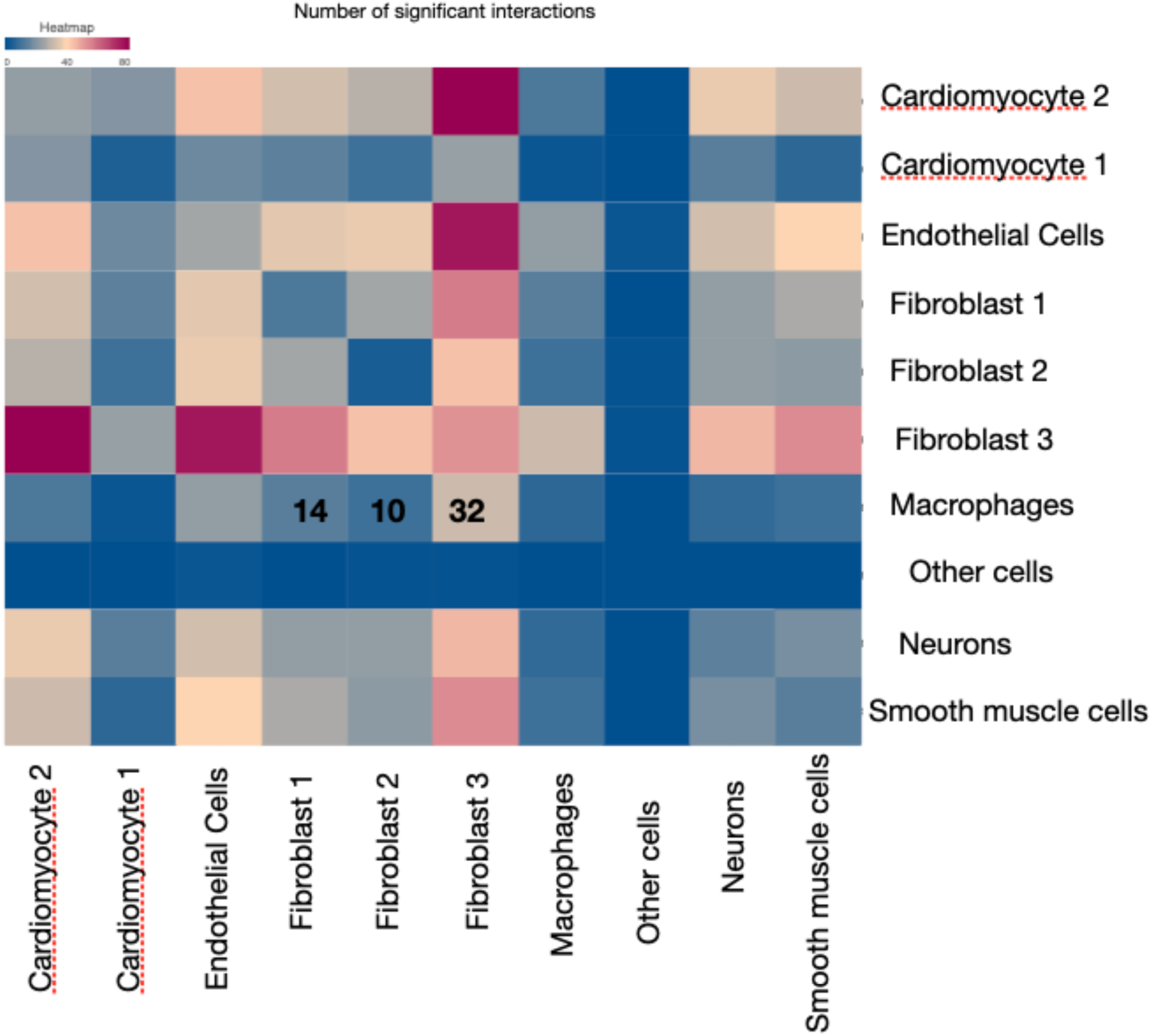
Myeloid-*Sting* regulates intercellular crosstalk in the heart after myocardial infarction. Cell communication analysis using Cellphone DB in the infarct of *LyzM^Cre^*^/+^ *Sting*^fl/fl^ mice show interactions among various cardiac cell populations.

**Figure S13:**
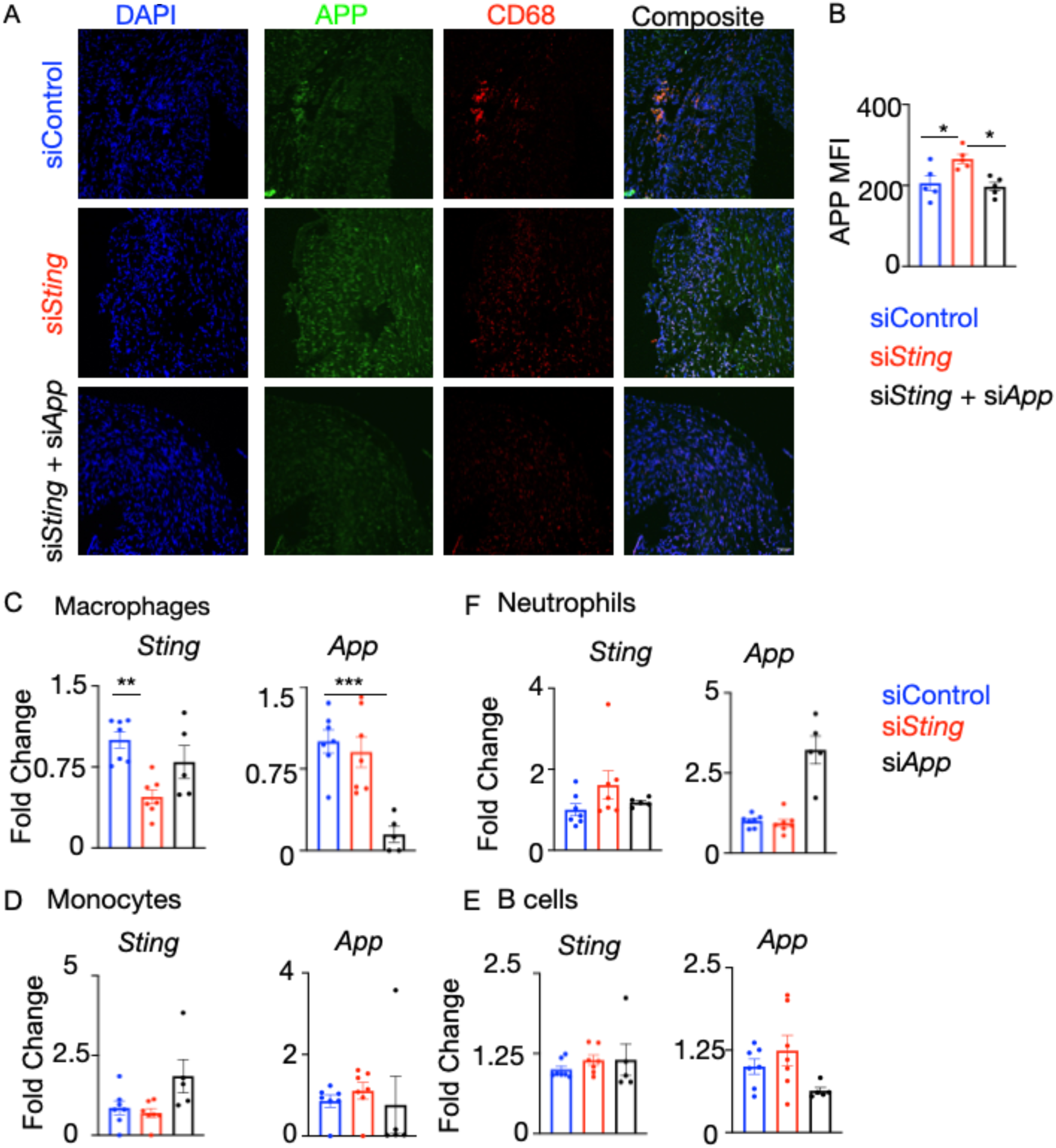
Macrophage Sting suppresses App. si*App* incorporated in lipidoid nanoparticles silenced *App* specifically in macrophages. C57BL/6 mice were injected intravenously with si*Control*, si*Sting,* or si*Sting+*si*App* i.v after MI. APP expression in left ventricle sections was quantified by immunostaining, representative sections (A), quantification (B), n=4 siSting, 5 siSting+siApp. *Sting* and *App* expression was measured in macrophages (C), monocytes (D), B cells (E), and neutrophils (F) by qPCR, n=5-7/group. The fold change in gene expression is in comparison to the siControl group. \**p*<0.05, \*\**p*<0.01, \*\*\**p*<0.001, one-way ANOVA with post-hoc Fisher LSD test.

**Figure S14:**
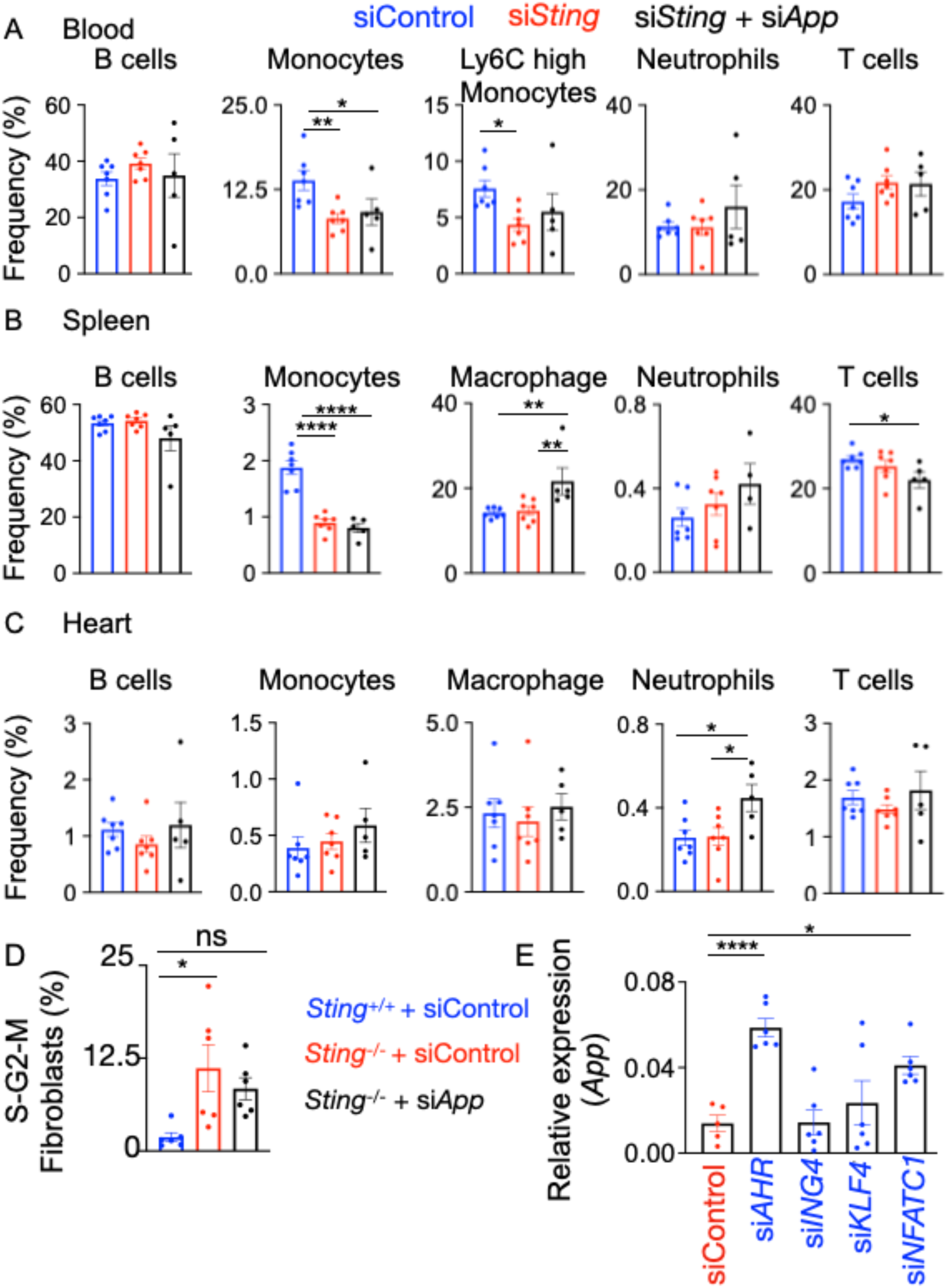
Macrophage *Sting* inhibits *App* to downregulate fibroblast proliferation. B cell, monocyte, Ly-6C^high^ monocyte, neutrophil, and T cell frequencies in the blood (A), spleen (B), and heart (C) of si*Control*, si*Sting,* and si*Sting+*si*App*-treated mice were evaluated by flow cytometry, n=6 *siControl*, *siSting*, n=5 *siApp*. D) Cell cycle progression of 3T3-L1 fibroblasts treated with conditioned medium collected from *LyzM*^+/+^ *Sting*^fl/fl^, *LyzM^Cre^*^/+^ *Sting*^fl/fl^ BMDM transfected with siControl or si*App* was examined, n=6/group. E) Expression of *App* in BMDM upon silencing key transcription factors regulated by *Sting* is shown, n=6/group. \**p*<0.05, \*\**p*<0.01, \*\*\**p*<0.001, \*\*\*\**p*<0.0001, one-way ANOVA with post-hoc Fisher LSD test.

**Figure S15:**
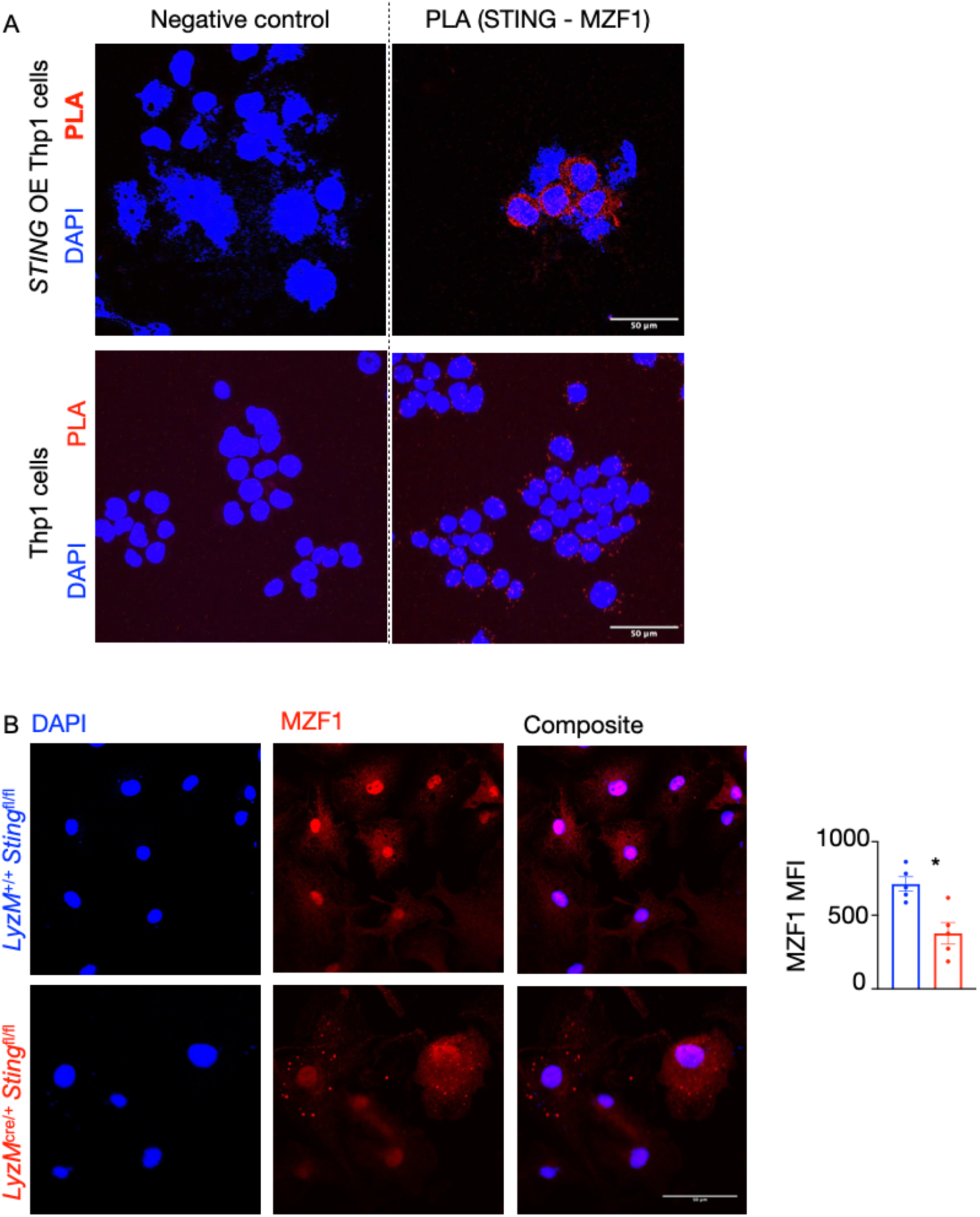
STING interacts with the transcription factor MZF1. A) Proximity ligation assay was performed to probe the interaction between STING and MZF1 in THP1 cells with (upper panels) or without (lower panels) STING overexpression. Scale bar: 10μm. B) MZF1 was quantified in BMDM obtained from *LyzM^+^*^/+^ *Sting*^fl/fl^ and *LyzM^Cre^*^/+^ *Sting*^fl/fl^ mice by confocal microscopy, n=4/group. \**p*<0.05, 2-tailed Student’s t-test.

## Methods

All animal experiments were approved by the University of Pittsburgh Institutional Animal Care and Use Committee (IACUC) and compliant with ethical guidelines and welfare.

### Human heart specimens

Deidentified human left ventricle specimens were obtained from the NDRI (IRB approval #STUDY19090164). Specimens were fixed, and embedded in OCT compound (Fisher Scientific # 23-730-571) for immunostaining and FFPE blocks for histological staining. OCT blocks were sectioned (10 µm), and sections were permeabilized with 0.1% Triton X-100 (Millipore Sigma # MTX15683) in PBS prior to staining, TOM20 (Proteintech # CL488-11802)), CD68 (Abcam # ab224029), 8OHdG (Novus bio # NB600-1508-50ul) or FEN1 (Thermo Fisher # MA1-23228), APEX2 (Thermo Fisher #BS-6587R)or pSTING (Thermo Fisher # PA-105674) antibodies. Sections were imaged with a Nikon A1 confocal microscope. Images were quantified in a randomized, blinded manner using ImageJ (NIH).

### Quantification of cell free mtDNA from human plasma

DNA was isolated from plasma samples obtained from human control subjects and MI patients using QIAamp MinElute Virus Spin Kit. (Qiagen,Cat# 57704), per manufacturer’s instructions. Mitochondrial DNA levels were quantified by quantitative PCR (qPCR) using human mitochondrial specific primers: h Mito F (5’-CACTTTCCACACAGACATCA-3’) and h Mito R (5’-TGGTTAGGCTGGTGTTAGGG-3’). For the absolute quantification, mt DNA was isolated from human heart tissue using mitochondrial DNA isolation kit (Abcam; cat#ab65321). Serial dilutions of the purified mtDNA were prepared, and a standard curve was generated by plotting qPCR Cq values (hMito) against mtDNA concentrations measured by NanoDrop spectrophotometry. Absolute mtDNA levels in plasma samples were calculated by extrapolating sample Cq values to the standard curve.

### Animals

All animal experiments were performed according to the NIH guidelines, and the protocols of the animal experiments were approved by the University of Pittsburgh Institutional Animal Care and Use Committee. The mice were housed in 12-hour dark/light cycle at 70° F and 40% humidity. Adult C57BL/6 wild-type (#000664), *LyzMCre*^/cre^ (#004781), and *Sting*^fl/fl^ (#031670) mice were obtained from The Jackson laboratory. Myeloid-*Sting* deficient mice were generated by breeding *Sting*^fl/fl^ mice with *LyzMCre*^/cre^ mice. MI was induced in 10-12 week old *LyzM*^+/+^ *Sting^fl^*^/fl^ and *LyzM*^Cre/+^ *Sting^fl^*^/fl^ animals by permanent ligation of the left anterior descending artery as previously described ^75^. Age matched litter mate controls were used in each experiment.

### In vivo gene silencing with lipidoid nanoparticles

MI was induced by permanent LAD ligation in 10-12-week old female C57BL/6 mice. si*Control* or siRNA (si*Polg, siPgc1a, siCmpk2, siSting, sicGAS, siApex2, siFen1,* or *siApp*) formulated in lipidoid nanoparticles (DOTAP liposomes) were administered i.v. twice a week for 8 weeks after MI. As previously described, 50:50 DOTAP:cholesterol liposomes (Encapsula Nanosciences # GEN-7001), were administered at 1:7.5 siRNA:total DOTAP by weight and a dosage of 1.5mg/ kg DOTAP as previously described ^32^.

### Organ harvesting and flow cytometry

Mice were euthanized according to the IACUC guidelines at the University of Pittsburgh. PBS was perfused through left ventricle, and hearts were harvested for either flow cytometry or histological analysis. For flow cytometry, hearts were minced, and digested in 450 U/ml collagenase I (sigma #C0130), 125U/ml collagenase XI (Sigma # C7657), 60 U/ml DNase I (Sigma # D5319), 60 U/ml hyaluronidase (Sigma #H3506), and 20 µM HEPES (Corning # 25-060-CI) at 37° C and 750 rpm for 20 minutes. The dissociated cell suspension was passed through 70 µM cell strainers and resuspended in FACS buffer (PBS+0.5%BSA (Sigma# A9647) after centrifugation. For histological analyses and scRNAseq, hearts were fixed in 10% neutral buffered formalin (Sigma # HT501128-4L) for 18 hours and embedded in FFPE blocks. Blood was collected by terminal cardiac puncture and incubated with RBC lysis buffer (BioLegend #420302) for 3 minutes at room temperature, followed by addition of FACS buffer and centrifugation to pellet leukocytes. A hemocytometer was used to count the number of viable cells in the organs.

The following panel of antibodies were used to analyze the myeloid cell population in mice: anti-CD45.2 (Biolegend 109820, clone 104, lot B364148), CD11b (BD Biosciences 557657, clone M1/70, lot 1271559), CD115 (Biolegend 565249, clone T38-320, lot 7291701), Ly6G (Biolegend 563979, clone 1A8, lot 0269936), Ly-6C (eBioscience 45-5932-82, clone HK1.4, lot 2162018), F4/80 (Biolegend 123114, clone BM8, lot B370041) CD3 (BD Biosciences 367-0032-82, clone 17A2, lot 2526889), and CD19 (BD Biosciences 563148, clone 1D3, lot 3130912). Neutrophils were identified as CD11b^+^, Ly6G^+^, and CD115^−^. Monocytes were considered as CD11b^+^, Ly-6G^-^, and CD115^+^. B and T lymphocytes were identified as CD11b^-^, CD19^+^, CD3^-^ and CD11b^-^, CD19^-^, CD3^+^ respectively. A Cytek Aurora flow cytometer was used to acquire the data, which were analyzed with FlowJo software (Tree Star).

### Histology, immunofluorescence, and western blotting

For histology and single-cell RNAseq, hearts were harvested after PBS perfusion and fixed in 10% neutral buffered formaldehyde. 50μm Sections were cut using a microtome for single-cell RNAseq analyses, and 5μm sections were cut for histology. Masson Trichrome staining was performed to quantify fibrosis as follows. Sections were fixed overnight in Bouin’s fixative (RT). Slides were washed in tap water to remove residual Bouin’s fixative, then stained for 10 minutes with a 1:1 mixture of hematoxylin A and B (Thermo Scientific). Next, the slides were incubated in scarlet acid stain (Electron Microscopy Sciences) for 10 minutes, followed by 5 minutes in phosphomolybdic acid solution (Electron Microscopy Sciences). Finally, the sections were stained with aniline blue solution (Electron Microscopy Sciences) for 5 minutes and then transferred to 1% acetic acid for 5 minutes, dehydrated in ethanol and xylene, and mounted in permount (Electron Microscopy Sciences), cured for 24 hours before imaging. For immunostaining, hearts were embedded in OCT compound. Stained sections were imaged using Nikon A1, a confocal microscope, at the University of Pittsburgh Center for Biological Imaging. ImageJ (NIH) was used for analyses.

### Luminex

Biomarkers were analyzed using a Bio-Rad mouse 23-plex and a Bioplex 200 (Bio-Rad). Serum samples were diluted 5-fold using diluent provided by the company and analyzed in duplicates. Cell culture supernatants were analyzed as neat and run in single. Standard curve and experimental data were generated and analyzed using the Bio-Plex Manager Software.

### RT-PCR

RNeasy RNA isolation kit (Qiagen # 74106) was used to extract RNA from BMDM, and cDNA was synthesized using Applied Biosystems High Capacity RNA to cDNA kit (#4374967). Gene expression was quantified by quantitative RT-PCR using SYBR green mastermix (Applied Biosystems #A25778, and primers (IDT). Gene expression Ct values were normalized to those of β-actin. Heat maps were generated with Excel.

### BMDM culture, and oxidized LDL and siRNA treatments

Bone marrow cells were isolated from the femurs and tibias of *LyzM*^+/+^ *Sting*^fl/fl^ and *LyzM*^Cre/+^ *Sting*^fl/fl^ mice after euthanasia followed by intracardiac perfusion using PBS. The cells were cultured in low-glucose DMEM containing 5.5 mM glucose and supplemented with 20% L929 conditioned medium for 7 days. BMDM were passaged into 12-well plates, treated with 100 µg/ml concentrations of native or oxidized LDL (Lee Bio #360-10-0.5) for 48 hours. For gene silencing, either control or si*RNA* against the gene of interest (Silencer Select, Thermo Fisher Scientific) was transfected with Lipofectamine RNAiMax (Thermo #13778-150) according to manufacturer’s recommendations. Efficiency of gene silencing was evaluated by qPCR.

### Chromatin Immunoprecipitation and qPCR

Chromatin immunoprecipitation was carried out as previously described ^76^. Briefly, murine bone marrow-derived macrophages were transfected with *siControl* or *siSting*. Cells were harvested and chromatin was immunoprecipitated with anti-MZF1 antibody. Following chromatin enrichment and DNA extraction, qPCR was performed with the primers listed below to determine relative enrichment of MZF1 on the APP primer region.

ChIP qPCR primers:

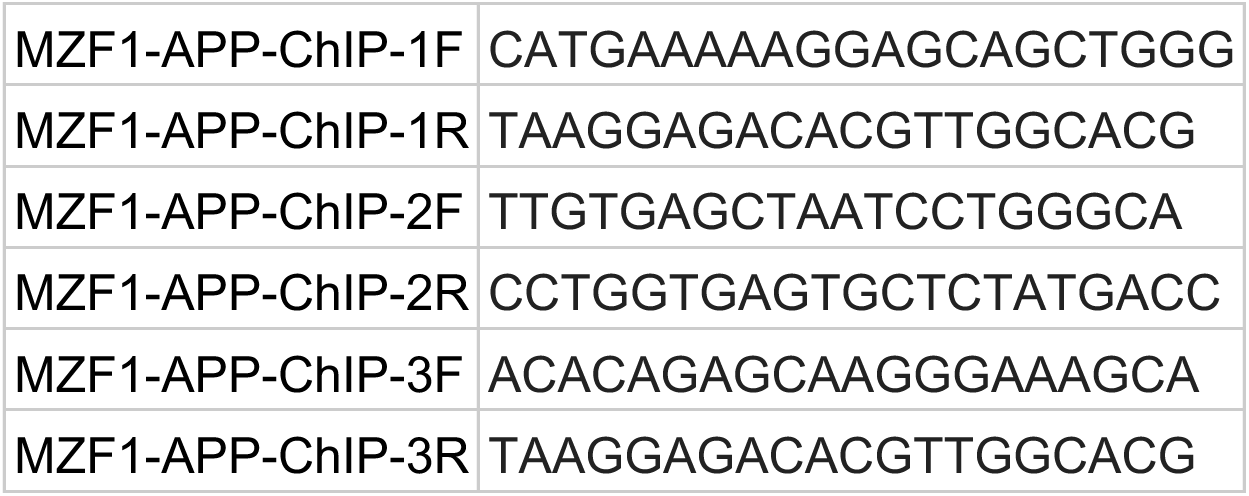

### Single-cell RNA sequencing

FLEX (Fixed RNA Profiling) library preparation using the 10x Genomics platform was performed on FFPE tissue sections from *LyzM*^+/+^ *Sting*^fl/fl^ and *LyzM*^Cre/+^ *Sting*^fl/fl^ hearts. Cells were isolated from 50 μm sections following xylene based deparaffinization and gentle MACS Octo dissociation. Cell yield and quality were assessed using NucSpot 470 staining on a Revvity Cellometer Ascend. Cells were stored at -80°C in long term storage buffer when not immediately processed. Fixed cells were hybridized with probe barcodes and partitioned into nanoliter scale Gel Beads-in-emulsion (GEM) droplets using a microfluidic chip. A pool of approximately 737,000 10x GEM barcodes was used to index the contents of each partition. Samples were equally pooled per GEM reaction using uniquely barcoded probe sets to generate a single library. Final libraries were assessed for size using an Agilent TapeStation. 49680 cells per sample were loaded for library preparation. Library concentrations were measured using an Invitrogen Qubit 2.0 fluorometer. Libraries were sequenced on an Illumina NovaSeq X Plus platform at 10000 reads/cell.

Count matrices (h5) generated from the FASTQ files were used for downstream analysis. Data were analyzed using Partek Flow software (v12.10.0). Single cell quality control was performed in Partek Flow by retaining cells with ≤ 12,000 read counts, ≤ 2,500 detected features, ≤ 8% mitochondrial reads, and ≤50% ribosomal reads. Filtered cells were then processed to exclude lowly expressed features, with expression values ≤ 1 in ≥ 99% of cells, followed by re-evaluation of the single cell QC matrices, annotation merging, and generation of expression distribution statistics in Partek Flow. Data were total count normalized, log_2_ transformed in accordance with Partek flow recommended best practices. Principal component analysis (PCA) was performed to compute the top 10 principal components, and cells were clustered using a graph-based clustering approach with Louvain algorithm. To visualize cellular heterogeneity, dimensionality reduction was performed using t-distributed stochastic neighbor embedding (t-SNE) and uniform manifold approximation and projection (UMAP). Cell type annotation was performed by comparing the top five marker genes from each cluster with reference signatures in the PangalaoDB database. Differential expression in macrophages was performed using MAST handle modeling with FDR correction, comparing *LyzM*^+/+^ *Sting*^fl/fl^ MI vs sham and *LyzM*^+/+^ *Sting*^fl/fl^ vs *LyzM*^Cre/+^ *Sting*^fl/fl^ MI at the transcript level. Differentially expressed features were filtered using an FDR step up threshold ≤ 0.05 for multiple comparison correction and log_2_ fold change cutoff excluding values between -2 and 2, generating a gene list for downstream analysis. Heatmaps were generated from the expression data using hierarchical clustering. The same differential expression and filtering workflow was applied to Fibroblast 1, Fibroblast 2 and Fibroblast 3. Pathogenic fibroblasts among the three fibroblast populations were identified by applying pathogenic fibroblast gene signatures from Kuppe *et al*.,Nature (2021) and Amrute *et al.,* Nature (2024) independently. Signature gene list was mapped onto cell cluster visualizations to assess expression patterns across fibroblast clusters, revealing fibroblast 2 as pathogenic subset. Cell to cell communication analysis was performed using CellPhoneDB. To further interrogate specific interactions, reports were generated for macrophages and fibroblast 2, summarizing ligand-receptor interactions for a curated gene set and interaction score and significant interaction outputs for the downstream visualization and interpretation.

Canonical and disease pathway analysis were performed on differentially expressed genes (DEGs) identified from single cell RNA sequencing of macrophages and fibroblast annotated from *LyzM*^+/+^ *Sting*^fl/fl^ Sham, MI, and *LyzM*^Cre/+^ *Sting*^fl/fl^ MI mice heart, using Qiagen Ingenuity Pathway Analysis (IPA; v153384343).

#### Proximity Ligation Assay

Proximity ligation assay (PLA) was performed by using the Duolink® In Situ Red Starter Kit Mouse/Rabbit (Sigma-Aldrich; DUO92101) according to the manufacturer’s instructions. PLA was conducted in both normal THP-1 cells and STING overexpressed THP-1 cells. Primary antibodies used were anti- MZF1 (Abcam; Cat#ab64866) anti STING (Invitrogen; MA5-26030). Single primary antibody incubations were included as negative controls. Fluorescent PLA signals were imaged using a Nikon A1confocal microscope.

### Statistical analysis

Data are displayed as mean ± SEM. Statistical significance between groups was performed using Graphpad PRISM with Student’s t-test or ANOVA according to the dataset. Results were considered as statistically significant when p < 0.05.

## Data Availability

The single-cell RNAseq data generated in this study have been deposited in GEO.

